# Unveiling the role of Tdark genes in genetic diseases and phenotypes through bioinformatics-based functional enrichment and network analyses

**DOI:** 10.1101/2024.01.13.575505

**Authors:** Doris Kafita, Kevin Dzobo, Panji Nkhoma, Musalula Sinkala

## Abstract

In recent years, there has been growing interest in understanding the role of dark genes in genetic diseases and phenotypes. Despite their lack of functional characterisation, dark genes account for a significant portion of the human genome and are believed to play a role in regulating gene expression and cellular processes. We investigated the role of dark genes in genetic diseases and phenotypes by conducting integrative network analyses and functional enrichment studies across multiple large-scale molecular datasets. Our investigation revealed a predominant association of both dark and light genes with psoriasis. Furthermore, we found that the transcription factors UBTF and NFE2L2 are potential regulators of both dark and light genes associated with tuberculosis. In contrast, the transcription factors SUZ12 and TP63 are potential regulators of both dark and light genes associated with interstitial cystitis. Further network analysis of dark genes, including *CALHM6, HCP5, PRRG4, DDX60L* and *RASA2,* revealed a notably high weighted degree of association with genetic diseases and phenotypes. Moreover, our analysis revealed that numerous genetic diseases and phenotypes, including psoriasis, pick disease, tuberculosis, ulcerative colitis, interstitial cystitis, and Crohn’s disease, exhibited shared gene linkages. Additionally, we conducted a protein-protein interaction analysis to reveal 16 dark genes that encode hub proteins, including *R3HDM2, RPUSD4, FASTKD5*, and *MRPL15,* that could play a role in many genetic diseases and phenotypes, and are widely expressed across body tissues. Our findings contribute to the understanding of the genetic basis of diseases and provide potential therapeutic targets for future research. Identifying dysregulated dark genes in disease states can lead to new strategies for prevention, diagnosis, and treatment, thus advancing our understanding of disease mechanisms.

## Introduction

Genetic diseases and phenotypes can manifest as monogenic, polygenic, or complex conditions resulting from a combination of genetic and environmental factors [1–4]. Numerous clinical phenotypes are linked to mutations in specific genes, and disruptions in the regulatory mechanisms governing gene expression can also contribute to these conditions [5–10]. Diseases stemming from alterations in the human genetic code pose a substantial burden, with recognised genetic diseases affecting over 5% of live births and more than two-thirds of miscarriages [11]. In addition to highly penetrant monogenic disorders and significant chromosomal changes, the heritability of common diseases strongly suggests a genetic basis for prevalent disorders, such as cardiovascular disease [11–14].

Recent advances in genomic sequencing technology have led to the discovery of thousands of genetic variants and genes associated with genetic diseases and phenotypes [15–17]. The implementation of high throughput next-generation sequencing techniques has enabled the swift sequencing of a vast number of complete genomes, resulting in an ever-growing wealth of genomic information that is becoming increasingly accessible [18–21]. However, over one-third of all protein-coding genes, often referred to as the “dark genome” or “Tdark genes”, whose functions are either poorly understood or completely unknown [22–26], have not been extensively studied, resulting in limited literature coverage and a significant challenge in understanding their biological significance [22]. Recent research has revealed that targeting the dark genome could be a promising approach for cancer treatment since it shares similar genetic and pharmacological dependencies with the light genome [27]. However, our understanding of the roles of dark genes in other genetic diseases and phenotypes remains limited.

To better understand the dark genome, the Illuminating the Druggable Genome (IDG) project was launched in 2014 to identify knowledge gaps in human genome-encoded proteins and to explore understudied proteins that may be potential drug targets [24–26]. This project aims to gather comprehensive information on protein families, their associations with diseases and drugs, and their structural and functional properties [24,25]. Currently, the IDG project has generated vast amounts of data on over 20, 000 protein-coding genes [24,25]. These datasets contain information about the dark genome or dark genes, which have been curated based on the extent to which genes are studied. Furthermore, several other biomedical projects have curated information on protein-protein interaction [28–32], gene dependencies [33], genes involved in disease [34,35], and gene expression and regulation in human tissue [36].

The available large-scale datasets compiled by various biomedical databases present new opportunities to apply integrative analyses that could shed light on the dark genome and discover the possible roles of dark genes in human diseases. Here, we integrated data on dark and light gene annotations together with curated protein-protein interaction information to examine the connection between light and dark genes and genetic diseases and phenotypes. Additionally, we investigated the potential biological processes and pathways in which dark genes might participate, as well as their tissue-specific expression patterns. Overall, we illuminated various aspects of the biological functions involving the dark genome and offered insights into its potential involvement in genetic diseases and phenotypes.

## Methods

The study protocol was approved by the University of Zambia’s Health Sciences Research Ethics Committee IRB00011000. The analyses conducted in this study utilised publicly available datasets collected by the IDG, GTEx and HPA projects, which were made accessible through their respective project databases. The methods employed in this study adhered to the pertinent policies, regulations, and guidelines established by the IDG, GTEx and HPA projects for the analysis of their datasets and the reporting of findings.

### Determination of dark and light gene association with genetic diseases

To investigate the association between dark and light genes and genetic diseases/phenotypes, we obtained a list of genetic diseases, disease genes, and their associations from Pharos (version 3.15.1) (https://pharos.nih.gov/). Pharos serves as an integrated web-based informatics platform for analysis of data aggregated by the Illuminating the Druggable Genome (IDG) Knowledge Management Centre from various databases, including UniProt, GWAS, and DisGeNET, among others [25]. Genes/proteins in Pharos are classified into four target development levels, encompassing light genes (Tbio, Tclin and Tchem) and dark genes (Tdark).

Tbio designates targets in the early stages of development, exhibiting a well-documented Mendelian disease phenotype in the Online Mendelian Inheritance in Man (OMIM) database, Gene Ontology (GO) leaf term annotations supported by experimental evidence, or satisfying at least two of the following three criteria: a fractional PubMed publications count exceeding five, three or more annotations from the National Centre for Biotechnology Information (NCBI) Gene Reference Into Function (RIF), or 50 or more commercial antibodies, as indicated by data accessible from the Antibodypedia database [24,26].

Conversely, Tchem pertains to targets with available chemical tools for investigation, possessing bioactivity information in databases, with varying potency criteria depending on protein type (e.g., kinases, GPCRs) [26,37]. Tclin refers to well-characterised genes linked to drug mechanisms of action (MoA) and have established clinical relevance [24,26]. In contrast, Tdark designates targets with minimal information available, failing to meet any of the criteria set for Tclin, Tchem or Tbio [23,24].

We analysed the distribution of genetic diseases and their associated genes across each target development level and generated a heatmap to display the association of the top 40 genetic diseases with dark and light genes.

### Enrichment analysis of dark and light genes associated with selected common diseases

To gain insight into the regulatory mechanisms that control the expression and function of these genes. We conducted enrichment analysis on both light and dark genes that were linked to the top ten diseases, respectively. To accomplish this, we utilised eXpression2Kinases (X2K), a web tool developed by the Ma’ayan Laboratory (available at https://maayanlab.cloud/X2K/), which performs gene set enrichment analysis using various transcription factor gene set libraries, including integrated target genes for transcription factors determined by ChIP-seq experiments (ChEA). In addition, X2K employs the Genes2Networks (G2N) algorithm to identify proteins that interact physically with the transcription factors and performs kinase enrichment analysis (KEA) on the list of identified transcription factors and proteins using gene set libraries from kinase-substrate interaction databases [38]. Transcription factors and kinases with a hypergeometric p-value of less than 0.05 were considered statistically significant.

### Construction of the diseasome bipartite network for dark genes and their associated genetic diseases and phenotypes

We utilised Gephi (version 0.9.7), a robust network visualisation software [39], to further evaluate the role of dark genes and how they contribute to disease development using a bipartite network representing the diseasome. We used the circle pack layout with nodes grouped according to their category to visualise the network. The gene-diseasome network was made up of 2,704 nodes and 5,639 undirected edges. The nodes in this network belong to two different categories: diseases and genes. A connection between a disease and a gene was established if the gene was known to play a role in the disease/phenotype. Additionally, we manually classified each disease or phenotype into one of the 17 categories based on the physiological system affected. Furthermore, using the gene-diseasome bipartite network as a starting point, we used multimode network projection to create two biologically relevant network projections: the gene-disease network (GDN) and the disease-gene network (DGN). The GDN was visualised with nodes sized based on their degree within the range of 10 to 80. Nodes were coloured according to their associated disease/phenotype category. The Force Atlas 3D layout with default settings was applied to generate the network. Furthermore, to eliminate unconnected nodes, the giant component was used as a filter. Within the GDN, the nodes represented diseases, and two diseases were linked if they shared at least one gene that was associated with both. This network projection was composed of 503 diseases, with 5,262 unique connections. Furthermore, we used the Circle Pack layout that grouped nodes based on their weighted degree to visualise the DGN. Node size was determined by the weighted degree, ranging from 10 to 80. In addition, nodes were colour-coded according to their betweenness centrality. The DGN was composed of nodes that represented genes, with connections between two nodes occurring if the corresponding genes were both implicated in the same disease. The size of each node was proportional to the number of diseases in which the gene was involved. This network projection encompassed 2,147 genes with 467,344 unique connections. Additionally, we utilised Gephi to detect smaller clusters, or modules, within the bipartite network of the diseasome. To accomplish this, we employed the Louvain method, a widely used approach for cluster identification in network analysis [40]. To determine the significance of nodes in the networks, we calculated several network centrality metrics, such as degree, closeness centrality, betweenness, and eccentricity.

### Identification of hub dark genes

Further network analyses of the disease-gene network allowed us to identify key genes involved in disease processes. Using a list of the 2,144 dark genes, we obtained known protein-protein interaction of these genes from the University of California, Santa Cruz (UCSC) [28], ChIP Enrichment Analysis (ChEA) [29], Kinase Enrichment Analysis (KEA) [30], Search Tool for the Retrieval of Interacting Genes/Proteins (STRING) [31] and the Biological General Repository for Interaction Datasets (BioGRID) [32] to create an interaction network of dark genes –the network was visualised in Cytoscape (http://apps.cytoscape.org/), it had 149 nodes, with 216 edges. To identify the hub dark genes in the network, we used the *CytoHubba* (http://apps.cytoscape.org/apps/cytohubba) plugin in Cytoscape. Considering that biological networks are heterogeneous, we used more than one method to identify essential proteins [41]. Therefore, we used three algorithms to reflect the status of the node in the entire network from different aspects. We then obtained the top 20 genes from the protein-protein network ranked by three different algorithms: Closeness, Degree, and Maximal Clique Centrality (MCC). Furthermore, to observe the intersections of the results produced by the three algorithms, a Venn network was generated using Evenn (http://www.ehbio.com/test/venn) to identify the commonly predicted significant hub genes. Finally, to demonstrate the association of the identified hub dark genes with genetic diseases and phenotypes, a diseasome bipartite subnetwork was constructed using Gephi software.

### Gene ontology analysis of the identified hub dark genes

To elucidate the molecular mechanisms and biological processes underlying hub dark genes, we analysed hub genes at the functional level. Gene ontology (GO) enrichment [42] and Reactome pathway analysis [43] were performed using the Enrichr web [44] tool (https://maayanlab.cloud/Enrichr/). Pathways and GO terms with a P-value < 0.05 were considered significant.

### Tissue expression of hub dark genes

To gain insight into the molecular mechanisms by which the prominent dark genes we identified are associated with genetic diseases, we utilised the Genotype-Tissue Expression (GTEx) [45] database (version 8) (https://gtexportal.org/home/). Specifically, to analyse the expression patterns of the 16 hub dark genes across different tissues, we created a heatmap using the GTEx data. This allowed us to visualise the expression levels of the genes in different tissues and identify any patterns or correlations between gene expression and disease development. Additionally, we searched for expression quantitative trait loci (eQTLs) within the dark genes to better understand how mutations within these genes may contribute to disease development. By identifying eQTLs associated with the Tdark hub genes and the particular tissues in which they are expressed, we can begin to understand the biological pathways and mechanisms involved in disease development. By combining the eQTL data with our knowledge of disease-gene associations, we hoped to gain a more comprehensive understanding of the molecular mechanisms underlying genetic diseases and identify potential therapeutic targets. Furthermore, we employed the Open Targets Genetics (https://genetics.opentargets.org), an open-access integrative resource that combines human GWAS and functional genomics data [46] to ascertain if any of the eQTLs are linked to phenotypes specific to tissues in which they are expressed.

### Tissue distribution and specificity analysis of dark and light gene expression

We performed the tissue distribution and specificity analysis of dark and light gene expression to gain insights into the functional characteristics and potential roles of these genes in different tissues. To achieve this, we obtained the gene expression data from the Human Protein Atlas (HPA) [47] (version 23.0) (https://www.proteinatlas.org/). This dataset provided us with information on tissue distribution and tissue specificity of gene expression.

HPA classifies genes based on tissue distribution and tissue specificity [48]. Tissue distribution is assessed based on the number of tissues with detectable mRNA levels above a specified cut-off and is classified into five distinct categories [48]: (a) Detected in single: only one tissue exhibits detectable levels. (b) Detected in some: more than one, but less than one-third of the tissues display detectable levels. (c) Detected in many: at least one-third, but not all tissues show detectable levels. (d) Detected in all: all 37 tissues exhibit detectable mRNA levels. (e) Not detected: None of the 37 tissues demonstrate detectable levels.

The classification of tissue specificity is determined by evaluating the fold-change in mRNA expression levels across 37 analysed tissues and organs, and it is categorised into five distinct specificity groups [48]: (a) Tissue enriched: signified by at least a fourfold higher mRNA level in one tissue compared to all other tissues. (b) Group enriched: characterised by a group of 2–5 tissues exhibiting at least a fourfold higher mRNA level compared to all other tissues. (c) Tissue enhanced: designated when one tissue demonstrates at least a fourfold higher mRNA level compared to the average level in all other tissues. (d) Low tissue specificity: identified when at least one tissue has mRNA levels above the cut-off, but the gene does not fit into any of the above categories. (e) Not detected: applicable when all tissues exhibit mRNA levels below the cut-off.

In addition, to determine whether the Xpresso model predictions are associated with tissue distribution and tissue specificity from HPA, we used the predictions from the Xpresso model, as presented by Agarwal and Shendure [49]. Furthermore, we used information from the Pharos database (version 3.15.1), including gene symbols and target development levels (Tbio, Tchem, Tclin and Tdark) for integration with tissue specificity and tissue distribution data.

### Data analysis

We utilised a combination of computational analyses, employing MATLAB R2021a and various bioinformatics tools, to conduct all analyses. Statistical significance was considered when p-values were < 0.05 for single comparisons or when the q-values (Benjamini-Hochberg adjusted p-values) were < 0.05 for multiple comparisons.

## Results

### Both dark and light genes are associated with genetic diseases and phenotypes

To investigate dark and light gene associations with genetic diseases and phenotypes, we utilised information from the Pharos [26] (version 3.15.1) web portal (https://pharos.nih.gov). Pharos provides gene classifications based on their target development levels (TDLs) (see Methods section for the description of TDLs). First, we evaluated the number of genes in each Pharos TDL category and found that most genes in Pharos belonged to the Tbio category (12,058, 59.07%), followed by Tchem (1,971, 9.67%), Tclin (704, 3.45%), and Tdark (5,679, 27.82%) (see S1a Fig).

We then integrated the information on TDLs with the genetic diseases and phenotypes (excluding cancers) associated with various genes. Here, our analysis revealed a significant variation in the number of genes and genetic traits across different TDLs. Specifically, Tbio genes comprised (7,642, 66.18%), Tdark genes (2,146, 18.58%), Tchem genes (1,327, 11.49%), and Tclin genes (432, 3.74%), as illustrated in Fig 1a. To view a comprehensive list of genes in each TDL, refer to S1 File.

**Fig 1.**
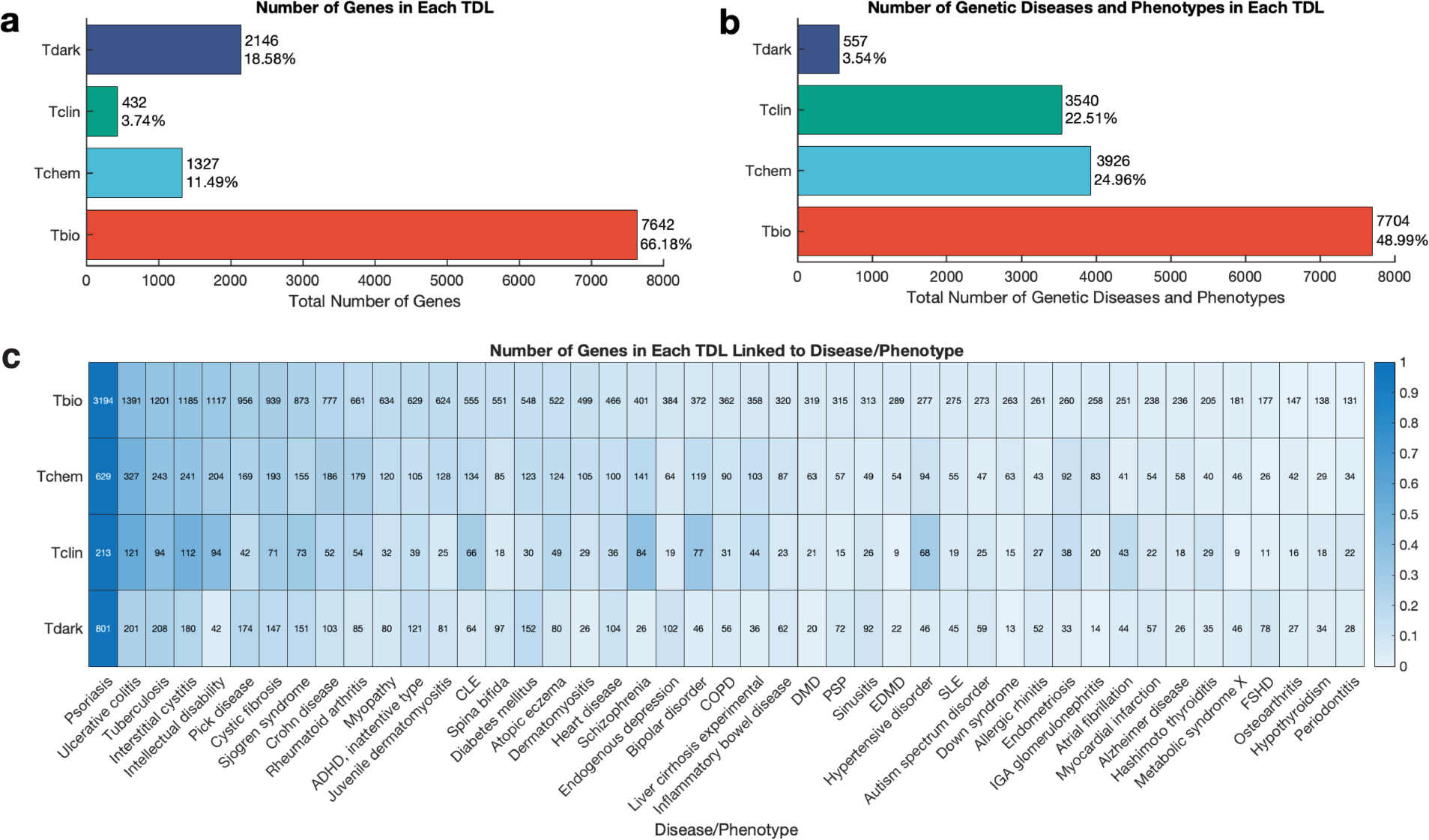
Distribution of genes and genetic diseases/phenotypes in each target development level (TDL). **a**. Frequency of genes. **b**. Frequency of genetic diseases and phenotypes. **c**. Distribution of genes linked to the top 40 genetic diseases and phenotypes. CLE: Cutaneous lupus erythematosus, COPD: Chronic obstructive pulmonary disease, DMD: Duchenne muscular dystrophy, PSP: Progressive supranuclear palsy, EDMD: Emery-Dreifuss muscular dystrophy, SLE: Systemic lupus erythematosus, FSHD: Fascioscapulohumeral muscular dystrophy.

Furthermore, we analysed the number of genetic diseases and phenotypes linked to genes within each TDL. We found that the Tbio category was associated with most diseases and phenotypes (7,704, 48.99% diseases and phenotypes), followed by Tchem (3,926, 24.96% diseases and phenotypes), Tclin (3,540, 22.51% diseases and phenotypes), and Tdark (557, 3.54% diseases and phenotypes), as shown in Fig 1b. The distribution of genetic diseases and phenotypes in each TDL is also available in S1 File.

Additionally, we observed that Tclin exhibited the highest average number of diseases per gene (8.194 diseases/gene), implying that these genes are associated with a broad spectrum of diseases. Conversely, Tbio and Tchem had lower averages (1.008 and 2.959 diseases/gene, respectively), indicating that these genes are less frequently implicated in disease. Furthermore, Tdark demonstrated the lowest average (0.259 diseases/gene), indicating that its genes are less comprehensively characterised (S1b Fig). Notably, Tdark exhibited a higher genes-to-disease ratio (3.853) compared to other TDLs, suggesting that, on average, there are approximately four genes associated with each disease in this category. This highlights the notable abundance of Tdark genes relative to identified diseases and underscores the need for continuous research to elucidate the roles of Tdark genes, potentially paving the way for innovative therapies across various diseases (S1c Fig).

In our analysis, we found that psoriasis had notable associations across the TDL categories: Tbio (3,194 associations), Tchem (629 associations), Tclin (213 associations), and Tdark (801 associations). Similarly, ulcerative colitis was significantly associated with Tbio (1,391 associations), Tchem (327 associations), Tclin (121 associations), and Tdark (201 associations). These were among the top genetic diseases and phenotypes associated with TDL gene categories (Fig 1c, also see S2 Fig). Interestingly, we found that some diseases, such as Galloway-Mowat syndrome 2 (X-linked), Bardet-Biedl syndrome 18 and chromosome 19q13.11 deletion syndrome, were exclusively associated with dark genes (see S1 File). Overall, these findings indicate a significant potential for uncovering the roles of Tdark genes in disease development, suggesting vast, untapped insights into their contributions to genetic disorders.

### Transcription factor and kinase enrichment analysis of dark and light genes associated with genetic diseases and phenotypes

To gain insight into the regulatory mechanisms involved in gene expression, we identified the top 10 genetic diseases and phenotypes with strong associations with both dark and light genes. These include conditions such as psoriasis, ulcerative colitis, intellectual disability, tuberculosis, and Crohn’s disease. Subsequently, we performed transcription factor and kinase enrichment analyses for genes in each TDL category, specifically targeting the top ten genetic traits, to better understand the regulatory mechanisms influencing gene expression. We first identified light genes associated with each disease and then assessed the enrichment of transcription factors and kinases specific to these genes. In parallel, we used the same method for the associated dark genes. This approach enabled us to delve into the distinct regulatory mechanisms for TDL gene categories in relation to each genetic trait (see Methods section).

Employing the in-silico Chromatin Immunoprecipitation Enrichment Analysis (ChEA) [29], we predicted significant enrichment (p < 0.05) of transcription factors potentially regulating the expression of light genes linked to each of the top ten genetic traits (refer to S2 File). Notably, rheumatoid arthritis and psoriasis stood out, with 36 and 35 predicted enriched transcription factors, respectively. The top-ranked transcription factors for light genes associated with psoriasis included SUZ12, which regulates 467 light genes (p = 1.5 x 10^-8^), SALL4, which regulates 130 light genes (p = 8.4 x 10^-8^) and GATA1, which regulates 235 light genes (p = 5.2 x 10^-6^) (Fig 2a). The top-ranked transcription factors for light genes associated with rheumatoid arthritis included RELA, which regulates 61 light genes (p = 2.7 x 10^-10^), UBTF, which regulates 134 light genes (p = 1.0 x 10^-8^) and GATA1, which regulates 77 light genes (p = 1.1 x 10^-7^) (S3a Fig, also see S2 File for a list of regulated genes from the input gene list).

**Fig 2.**
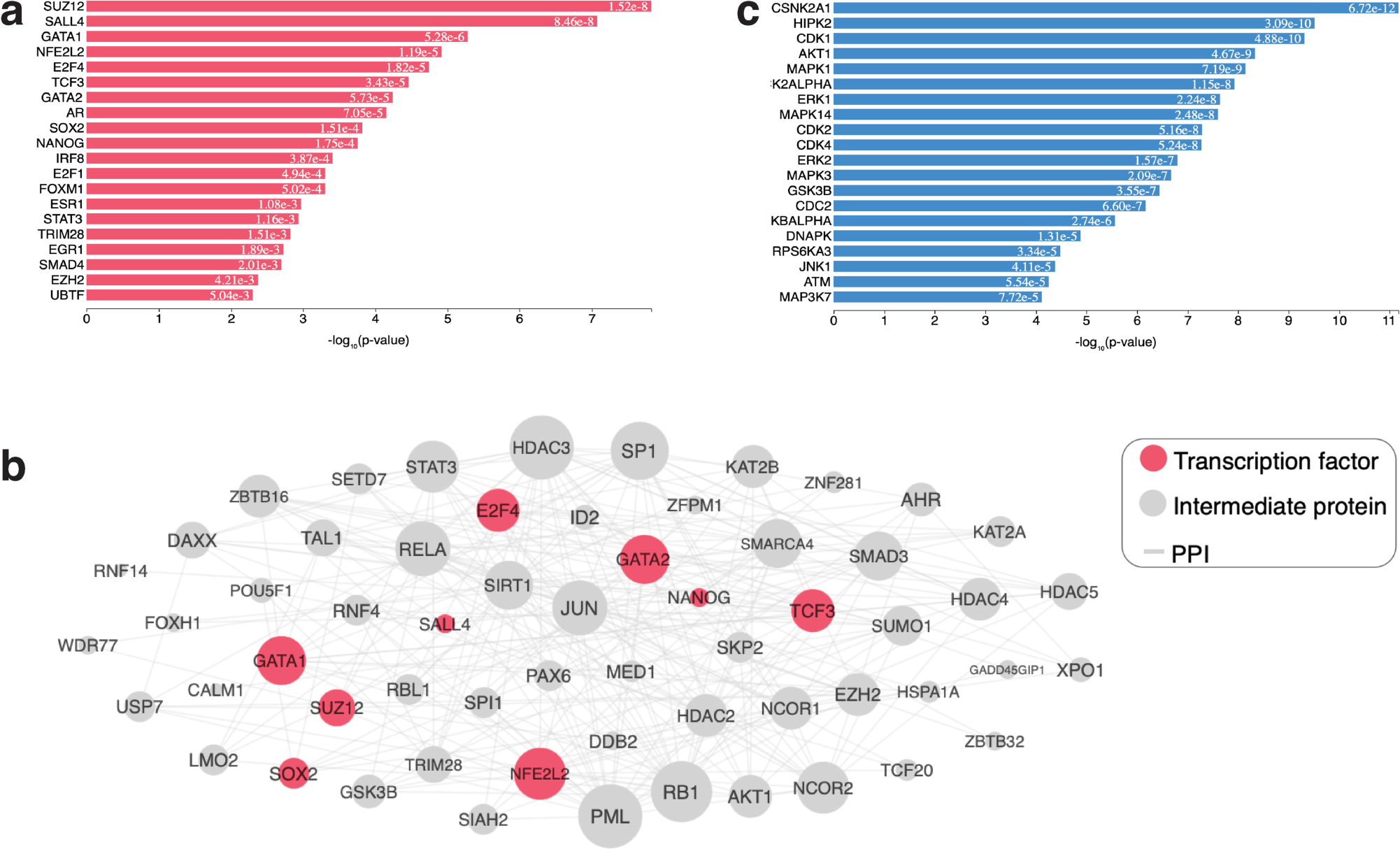
ChEA and KEA analyses of light genes associated with psoriasis. **a**. The top-20 predicted regulatory transcription factors. **b.** A subnetwork of connected transcription factors and their interacting proteins: the sub-network has 60 nodes with 526 edges. Transcription factors are pink nodes, whereas the proteins that connect them are grey. **c.** The top-20 predicted regulatory kinases.

To pinpoint the regulatory kinases for psoriasis and rheumatoid arthritis, we created a protein-protein interaction (PPI) subnetwork that maps the transcription factors and their interacting proteins (refer to Fig 2b and S3b Fig, respectively). We then employed the Kinase Enrichment Analysis (KEA) [30] to link the proteins in the PPI subnetwork to the protein kinases that likely phosphorylate them. Here, our result yielded a ranked list of protein kinases that likely regulate the transcriptome signature of psoriasis and rheumatoid arthritis. For psoriasis, among the top-ten ranked kinases were CSNK2A1 (p = 6.7 x 10^-12^), HIPK2 (p = 3.0 x 10^-10^), CDK1 (p = 4.8 x 10^-10^), and AKT1 (p = 4.6 x 10^-9^; Fig 2c, also see S2 File). Among the top-ten ranked kinases for rheumatoid arthritis were MAPK1 (p = 1.4 x 10^-13^), CK2ALPHA (p = 4.7 x 10^-13^), CSNK2A1 (p = 4.8 x 10^-13^), and MAPK3 (p = 2.0 x 10^-12^; S3c Fig, also see S2 File).

Conversely, focusing on the dark genes associated with the top ten genetic traits, we found significant transcription factors predicted in select cases through the ChEA [29] analysis, including ulcerative colitis, tuberculosis, interstitial cystitis, pick disease, diabetes mellitus, Sjogren syndrome, and cystic fibrosis (p < 0.05), as presented in S2 File. Furthermore, our analysis identified that among the dark genes associated with various genetic diseases and phenotypes, those linked to pick disease and tuberculosis displayed the most substantial enrichment, each being associated with ten transcription factors. This contrast underscores the differential regulatory landscape between dark genes across different diseases, with pick disease and tuberculosis exhibiting the most pronounced transcription factor enrichment. The top-ranked transcription factors for dark genes associated with tuberculosis included ELF1, which regulates 41 dark genes (p = 1.4 x 10^-3^); TCF7L2, which regulates 14 dark genes (p = 3.3 x 10^-3^) and UBTF, which regulates 28 dark genes (p = 5.5 x 10^-3^) (Fig 3a).

**Fig 3.**
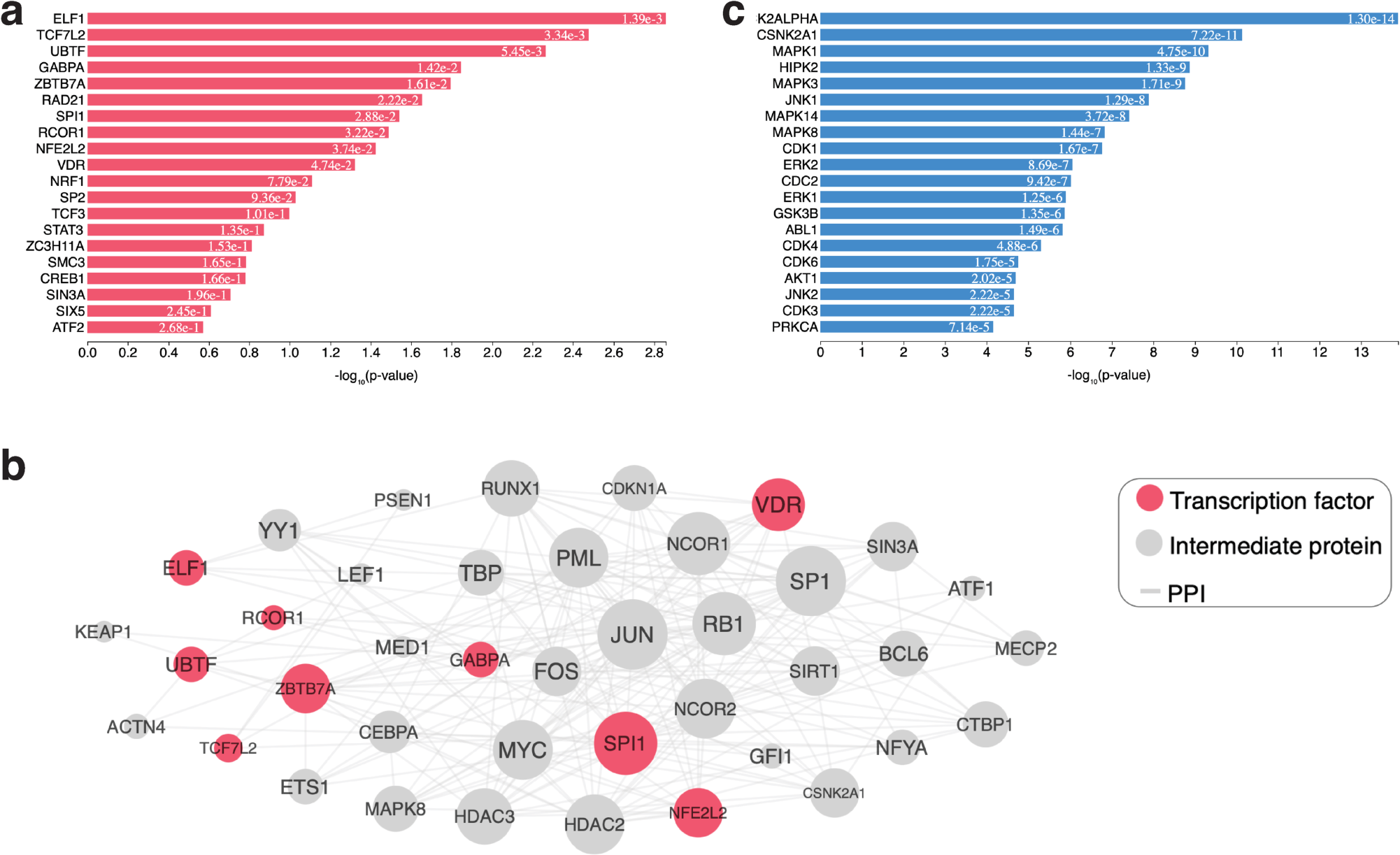
ChEA and KEA analyses of dark genes associated with tuberculosis. **a**. The top-20 predicted regulatory transcription factors. **b.** A subnetwork of connected transcription factors and their interacting proteins: the sub-network has 41 nodes with 393 edges. Transcription factors are pink nodes, whereas the proteins that connect them are grey. **c.** The top-20 predicted regulatory kinases.

The top-ranked transcription factors for dark genes associated with pick disease included TCF7L2, which regulates 12 dark genes (p = 4.4 x 10^-3^); BRCA1, which regulates 39 dark genes (p = 1.0 x 10^-2^); and PBX3 which regulates 19 dark genes (p = 1.1 x 10^-2^) (S4a Fig, also see S2 File for a list of regulated genes from the input gene list).

To identify the regulatory kinases of tuberculosis and pick disease, we created a PPI subnetwork of connected transcription factors and their interacting proteins (see Fig 3b and S4b Fig respectively). As previously, we utilised KEA [30] to link the proteins in the PPI subnetwork to the protein kinases that likely phosphorylate them. Here, our result yielded a ranked list of protein kinases that likely regulate the transcriptome signature of tuberculosis and pick disease. Among the top-ten ranked kinases for tuberculosis were CK2ALPHA (p = 1.3 x 10^-14^), CSNK2A1 (p = 7.2 x 10^-11^), MAPK1 (p = 4.7 x 10^-10^), and HIPK2 (p = 1.3 x 10^-9^; Fig 3c, also see S2 File). Additionally, the top-ten ranked kinases for pick disease were CSNK2A1 (p = 2.4 x 10^-12^), MAPK1 (p = 3.2 x 10^-12^), CDK1 (p = 2.3 x 10^-11^), and CDK4 (p = 1.9 x 10^-10^; S4c Fig, also see S2 File).

Interestingly, we discovered that several transcription factors potentially regulating gene expression of both light and dark genes in genetic diseases are shared. For instance, UBTF and NFE2L2 were identified as potential regulators of both the dark genes and light genes (S5 Fig) linked to tuberculosis. Similarly, we found that SUZ12 and TP63 were potential regulators of both the light and dark genes related to interstitial cystitis (see S2 File).

### Analysis of the diseasome bipartite network of dark genes and the associated diseases and phenotypes

To further shed light on the role of dark genes on disease development, we utilised the disease/phenotype-gene associations to construct a diseasome bipartite network (see Methods section). The bipartite network encompassed 557 genetic traits and 2,147 dark genes, of which 1,233 dark genes were linked to multiple diseases and phenotypes. The top-ranking dark genes associated with the most genetic traits were *TSEN34,* with 31 associations, followed by *PLPPR1,* with 29, and *LRMDA,* with 21. Furthermore, we found that most genetic traits in our diseasome were associated with the nervous system (Fig 4).

**Fig 4.**
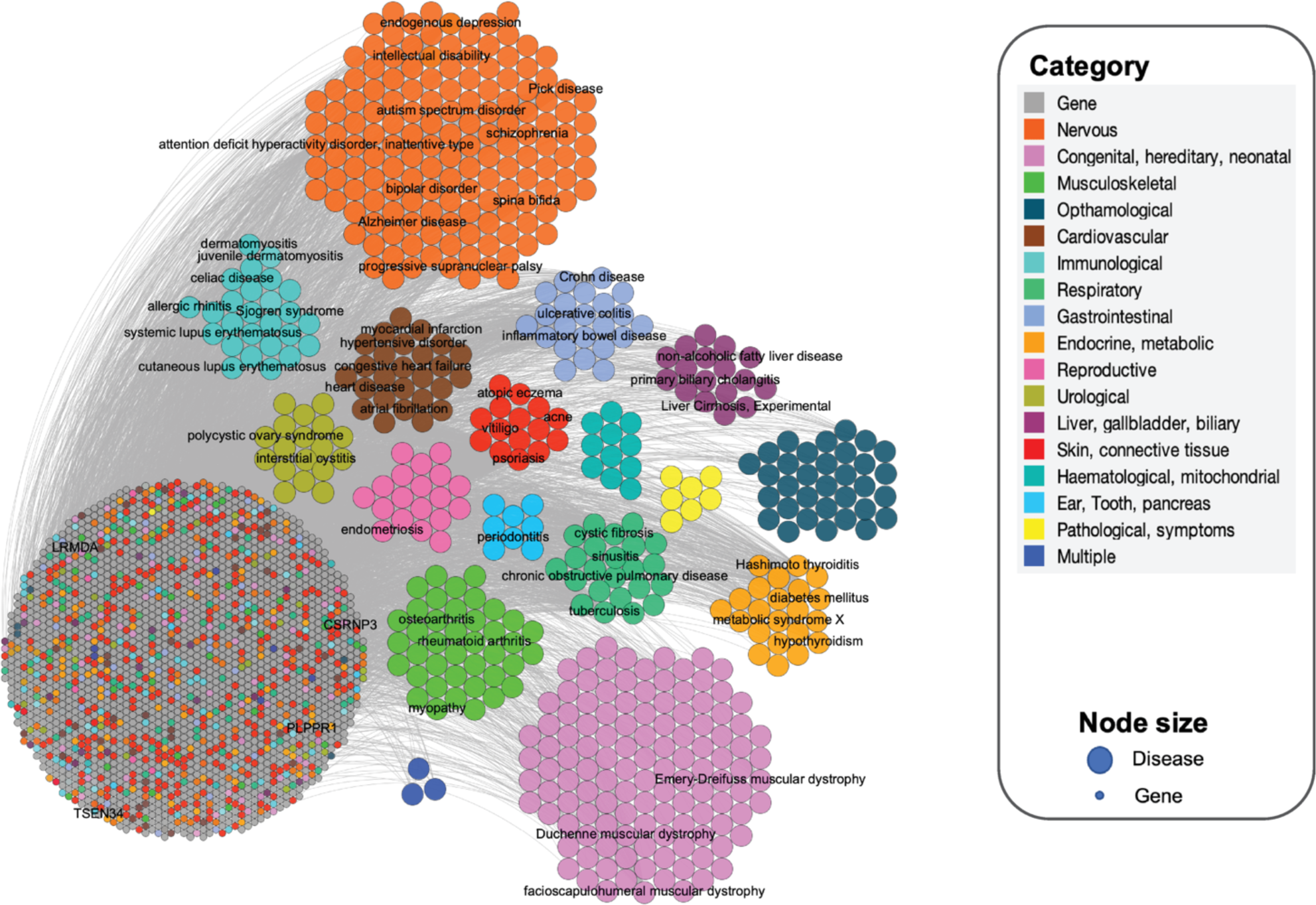
Gene-diseasome bipartite network. The bipartite network is composed of two disjoint sets of nodes with different sizes: the larger ones correspond to genetic diseases, whereas the smaller ones correspond to all genes. A link occurs between a disease and a gene if the gene is linked with the disease/phenotype. There are 18 categories in the diseasome bipartite network, as labelled in the legend. The links between disease pairs are shown in grey colour. Nodes are shaded grey if the corresponding genes are associated with more than one disease type. Disease/phenotype-specific gene nodes are coloured according to each disease/phenotype category, with “skin, connective tissue” having the highest number of disease/phenotype-specific genes. Similarly, disease/phenotype nodes are coloured based on the disease/phenotype category to which they belong, and most genetic diseases/phenotypes belong to the nervous system category. The names of diseases and genes with more than 20 connections were labelled in the network.

In addition, we employed multimode network projection on the gene-diseasome bipartite network to create two biologically meaningful network projections: the gene-disease network (GDN) for a disease-centred perspective (Fig 5) and the disease-gene network (DGN) for a gene-centred view of the gene-diseasome (Fig 6). Next, we analysed the GDN by calculating various network statistics including the weighted degree centrality, betweenness centrality and closeness centrality and eccentricity (see S3 File). Our analyses spotlighted psoriasis as the central hub of the network, demonstrating strong associations with most dark genes. Moreover, our degree distribution analysis indicated that most genetic traits (503 in total), such as psoriasis, pick disease, tuberculosis, ulcerative colitis, interstitial cystitis, and Crohn’s disease, exhibited shared gene associations across multiple diseases and phenotypes within the network. Our observation underscores the complex genetic interplay underlying various diseases and phenotypes (Fig 5, also see S3 File).

**Fig 5.**
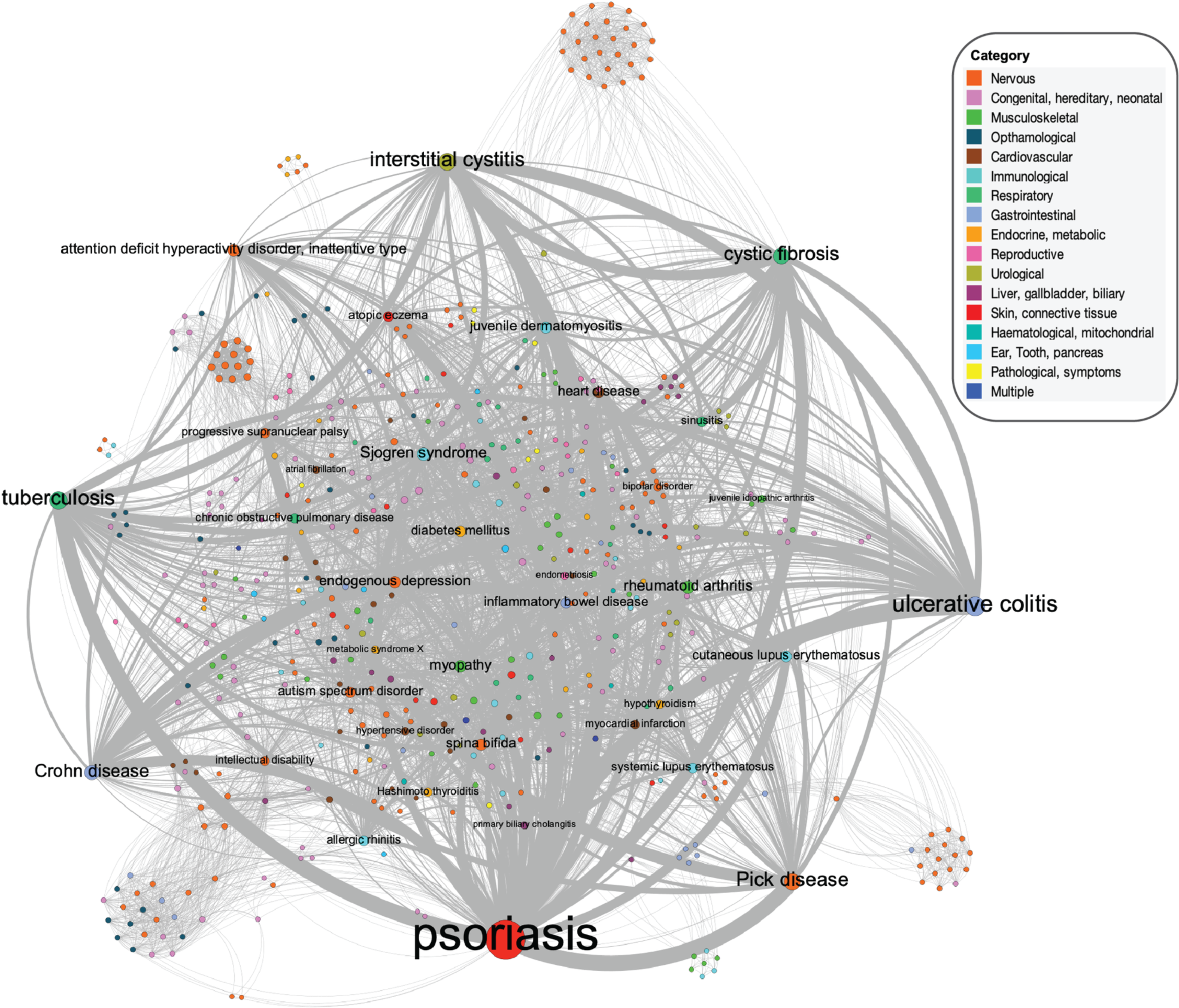
Gene-disease network (GDN). The GDN is the projection of the gene-diseasome bipartite network, in which nodes correspond to diseases/phenotypes, and two diseases/phenotypes are connected if there is at least one gene that is linked to both. The width of a link is proportional to the number of genes that are linked to both diseases. The size of a node is proportional to the number of genes linked to that disease. Different node colours are associated with different disease categories. The names of diseases with > 20 associated genes are labelled in the network. There are 17 disease categories in the gene-disease network (GDN), as labelled in the legend. The links between disease pairs are shown in grey colour. The weight of a link is proportional to the number of genes implicated in both diseases.

**Fig 6.**
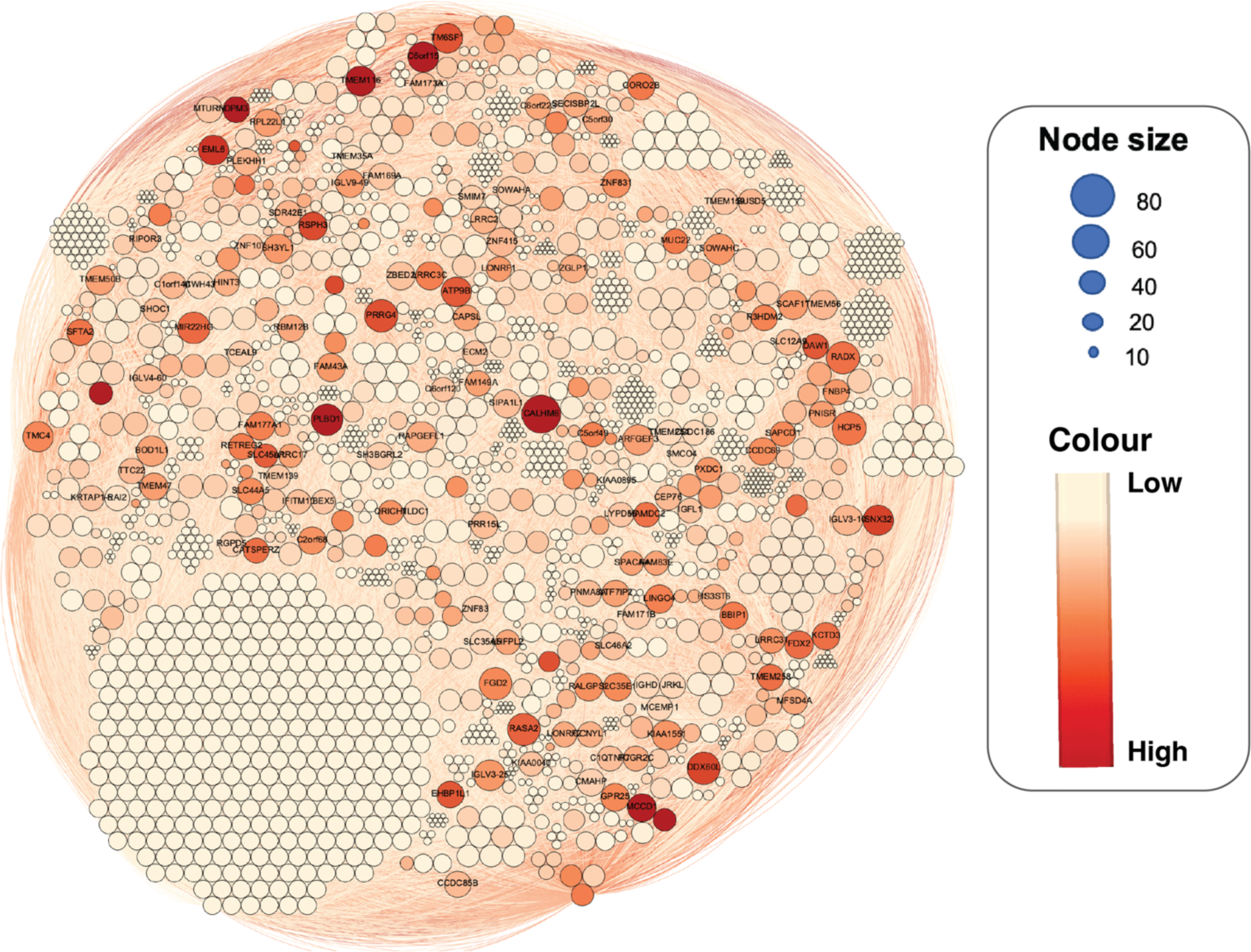
Disease-gene network (DGN). In the DGN, each node is a gene, with two genes being connected if they are implicated in the same disease. The size of the nodes was based on the average weighted degree (the larger the size, the higher the degree). The node colour exhibits betweenness centrality (the darker colour indicates the higher betweenness), Light orange is low, and red is high. The names of genes with a weighted degree range > 1119.0 are labelled in the network.

Furthermore, we calculated the network statistics on the DGN, including the weighted degree centrality, betweenness centrality, closeness centrality and eccentricity (see S3 File), which enabled us to identify dark genes with notably high weighted degree centrality, including *CALHM6, HCP5, PRRG4, DDX60L* and *RASA2*. These genes potentially play pivotal roles within the network, exerting strong influence over various genetic traits [50]. Furthermore, we pinpointed dark genes with high betweenness centrality, including *PLBD1*, *CTRB2*, *C6orf15*, *DPM3*, and *CALHM6*. These genes act as crucial connectors within the network, enhancing the flow of genetic information and possibly influencing the regulation of genetic traits [51,52]. This analysis deepens our understanding of the genetic foundations of complex traits and may pave the way for further research (Fig 6, also see S3 File).

Additionally, we identified seven smaller network clusters within the diseasome bipartite network and calculated a modularity score of 0.429 for these clusters. This positive score indicates the presence of a modularity structure [53,54] and is an average value for this network. This observation implies that genetic traits and their associated dark genes do not interact randomly but tend to cluster together in specific clusters. This finding provides further support for the notion that diseases sharing common genetic or biological foundations tend to cluster together, suggesting potential shared pathways or therapeutic targets [55–57]. Notably, modularity clusters in groups 2, 3, and 6 were the largest, representing distinct hub clusters with unique characteristics. The average degree for modularity class 2 was calculated as 3.064. The genetic traits heart disease, attention deficit hyperactivity disorder (inattentive type), and endogenous depression were highly connected with 82, 81 and 80 connections, respectively. Additionally, disease frequency in modularity class 2 showed that mostly nervous followed by musculoskeletal diseases belonged to this class. Gene *JRK* was found to have the highest degree, with 17 connections to various diseases (Fig 7; S3 File). The average degree for modularity class 3 was calculated as 3.975. The genetic traits ulcerative colitis, tuberculosis, interstitial cystitis, pick disease and Sjogren syndrome were highly connected with 159, 144, 135, 128, and 120 connections, respectively. Disease frequency in modularity class 3 indicated that mostly congenital, hereditary, neonatal, followed by nervous disease categories were in this group. Gene *C6orf15* was found to have the highest degree, with 14 connections to various diseases (Fig 7; S3 File). Furthermore, the average degree for modularity class 6 was calculated as 2.373. The genetic traits psoriasis and diabetes mellitus were highly connected with 409 and 88 connections, respectively. Disease frequency in modularity class 6 indicates that mostly nervous followed by congenital, hereditary, and neonatal disease categories were in this group. Gene *PLPPR1* was found to have the highest degree, with 28 connections to various diseases (Fig 7; S3 File).

**Fig 7.**
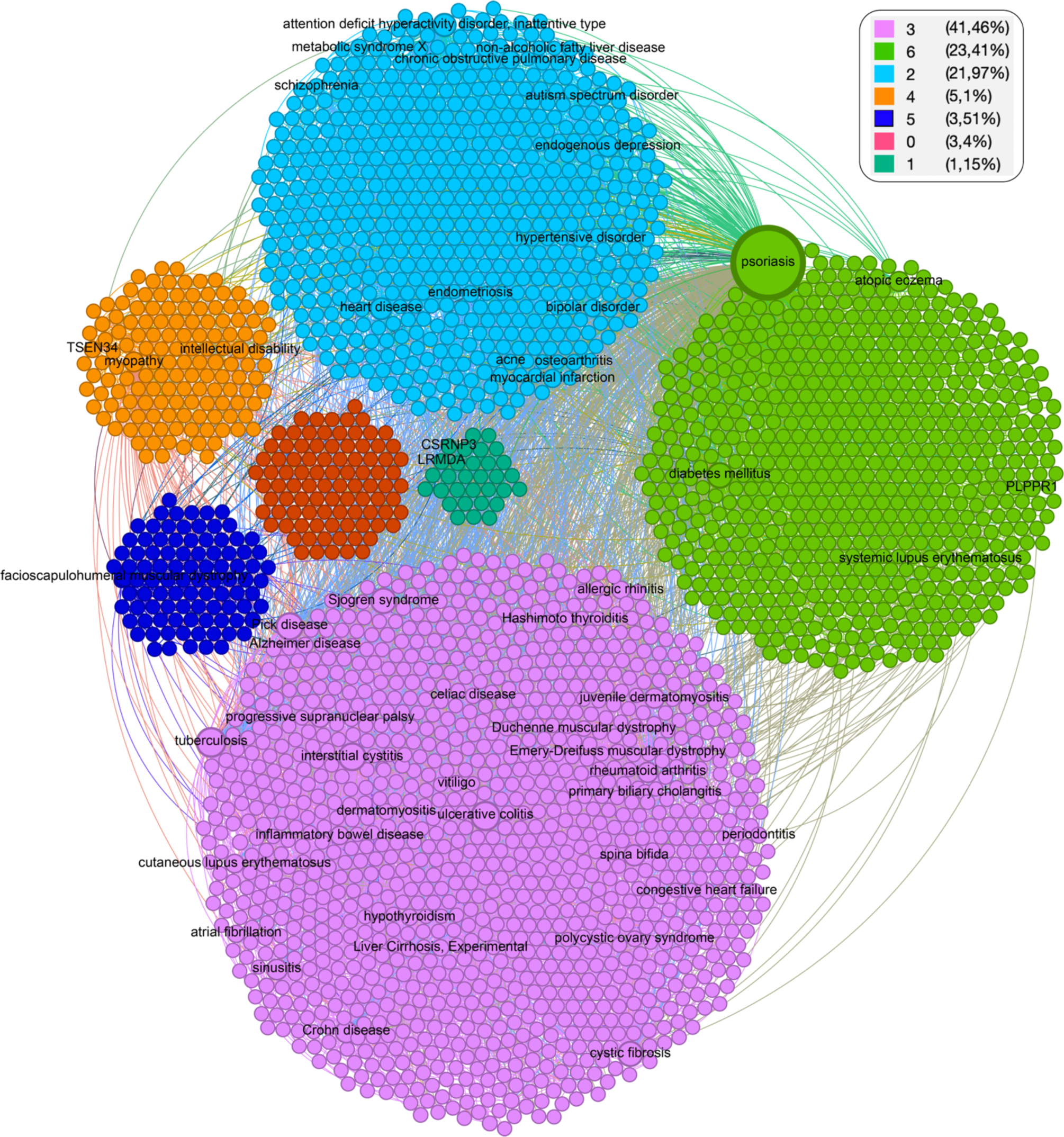
Cluster identification. The gene-diseasome bipartite network partitioned into seven distinct clusters, each highlighting a group of closely interconnected diseases and their corresponding genes that exhibit similar patterns of connections and associations within the network. The name of diseases and genes with > 20 connections are labelled in the network.

Overall, these findings indicate different levels of interconnectivity between the clusters in the network. This suggests that genetic traits within the same cluster may share similarities in terms of genetic factors and the potential underlying mechanisms [55–57]. We also noted variations in centrality metrics, including degree, betweenness, and closeness centrality, across the clusters, with certain clusters exhibiting higher centrality values compared to others (S3 File). Furthermore, the clear structural boundaries between clusters indicate that these clusters represent distinct subgroups within the network. Understanding the interconnectivity and centrality can assist in identifying key diseases and genes that are crucial in the context of disease interactions, pathways, and potential therapeutic targets [56,58–60].

### Identification of hub dark genes

We aimed to identify the key dark genes involved in disease processes. Out of the 2,147 dark genes we analysed, 1,998 dark genes could not be mapped to STRING [31], UCSC [28], ChEA [29], KEA [30], and BioGRID [32] databases and were therefore excluded from further analysis (see Methods section). The inability to map 1,998 of the dark genes to established databases underscores the existing gaps in understanding and the sparse representation of these genes within contemporary scientific literature and databases.

Subsequently, we visualised the PPI network using Cytoscape [61] [version 3.8.0]. The resulting PPI network of dark genes consisted of 149 nodes and 216 edges (Fig 8). The successful construction of the PPI network for the remaining dark genes suggests that despite their obscurity, these genes are not isolated entities and may play interconnected roles in biological systems.

**Fig 8.**
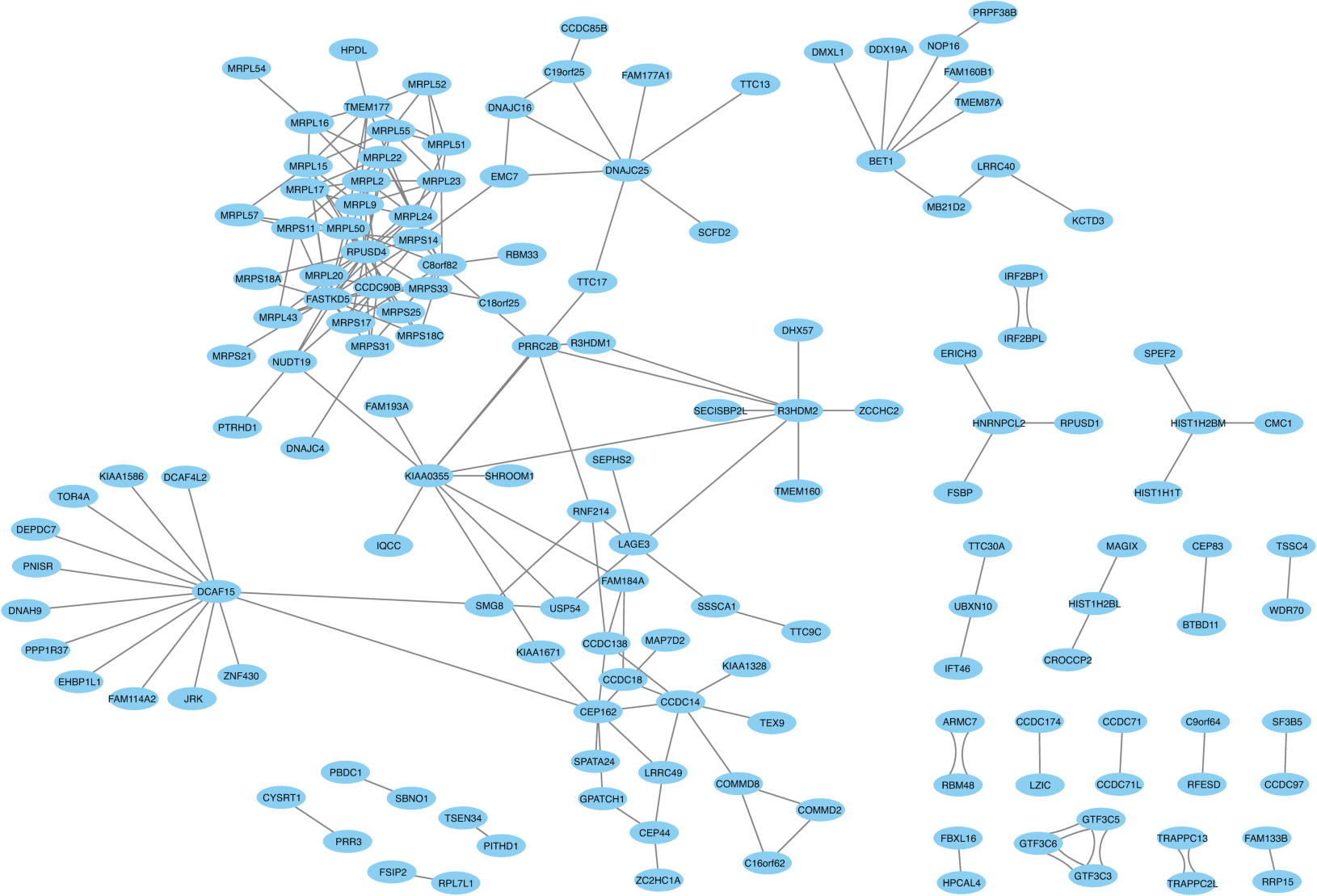
The protein–protein interactions network of dark genes was visualised in Cytoscape. The nodes indicate proteins, and the edges indicate the number of interactions. The PPI network had 149 nodes and 216 edges.

Moreover, we employed the *cytoHubba* [41] plugin to screen for the top 20 hub dark genes in the PPI network using three approaches: Maximal Clique Centrality (MCC), Degree and Closeness (see Methods section and S6 Fig). A total of 16 of the most significant genes emerged from the overlap of the results obtained from the three algorithms. Noteworthy among them were *RPUSD4, FASTKD5, MRPL15, MRPL9, DCAF15, R3HDM2, and CEP162* (S6d Fig). Notably, *R3HDM2* was the most interconnected, associating with nine genetic traits, including heart disease, psoriasis, and juvenile arthritis. Furthermore, *MRPL9* was linked with six traits, including myocardial infarction, allergic rhinitis, and inflammatory bowel disease (Fig 9).

**Fig 9.**
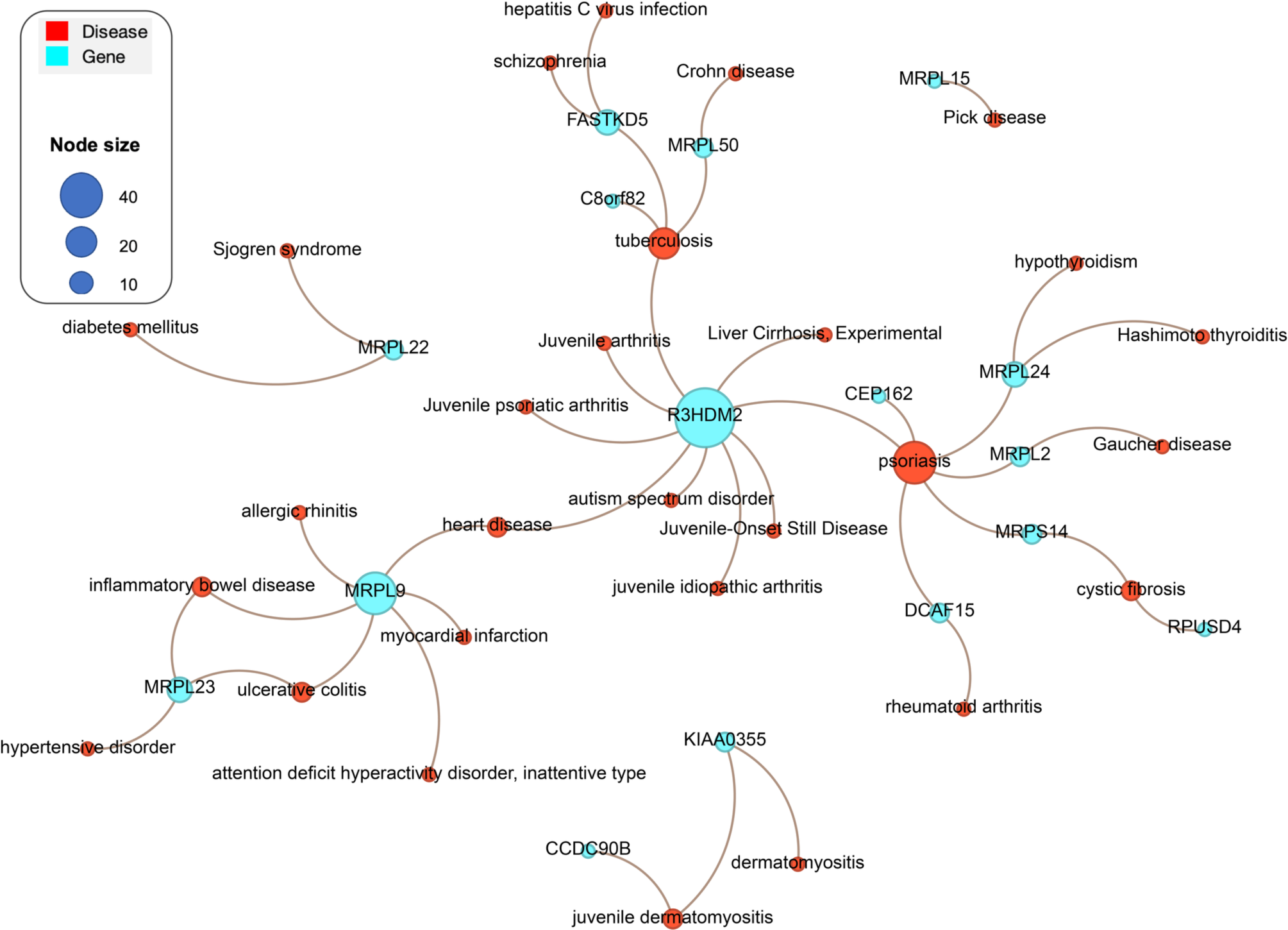
Diseasome bipartite subnetwork. Showing the interaction of genetic diseases with the identified hub dark genes. Node size represents degree, and node colour shows category (diseases coloured red and genes coloured blue).

### Function enrichment of hub dark genes

The high interaction of hub genes and their association with various diseases and phenotypes indicates that they may play important roles in biological processes. Therefore, to gain insights into the potential biological functions and pathways in which these genes may be involved, we performed functional enrichment analysis using Enrichr [44] (see Methods section). Our Gene ontology (GO) analysis showed that the hub genes are highly significantly enhanced for biological processes, the top enriched terms including mitochondrial translational elongation (p = 1.39 x 10^-15^) and mitochondrial translational termination (p = 1.39 x 10^-15^), (Fig 10a); Molecular functions including RNA binding (p = 2.97 x 10^-7^), and large ribosomal subunit rRNA binding (p = 3.99 x 10^-3^), (Fig 10b); Cellular components including mitochondrial inner membrane (p = 5.51 x 10^-11^), and organelle inner membrane (p = 8.44 x 10^-11^), (Fig 10c). As for biological pathways, the Reactome pathway analysis showed that hub genes are enriched with mitochondrial translation termination (p = 3.05 x 10^-11^), mitochondrial translation elongation (p = 3.05 x 10^-11^), and mitochondrial translation initiation (p = 3.05 x 10^-11^), (Fig 10d). The detailed results are shown in S4 File.

**Fig 10.**
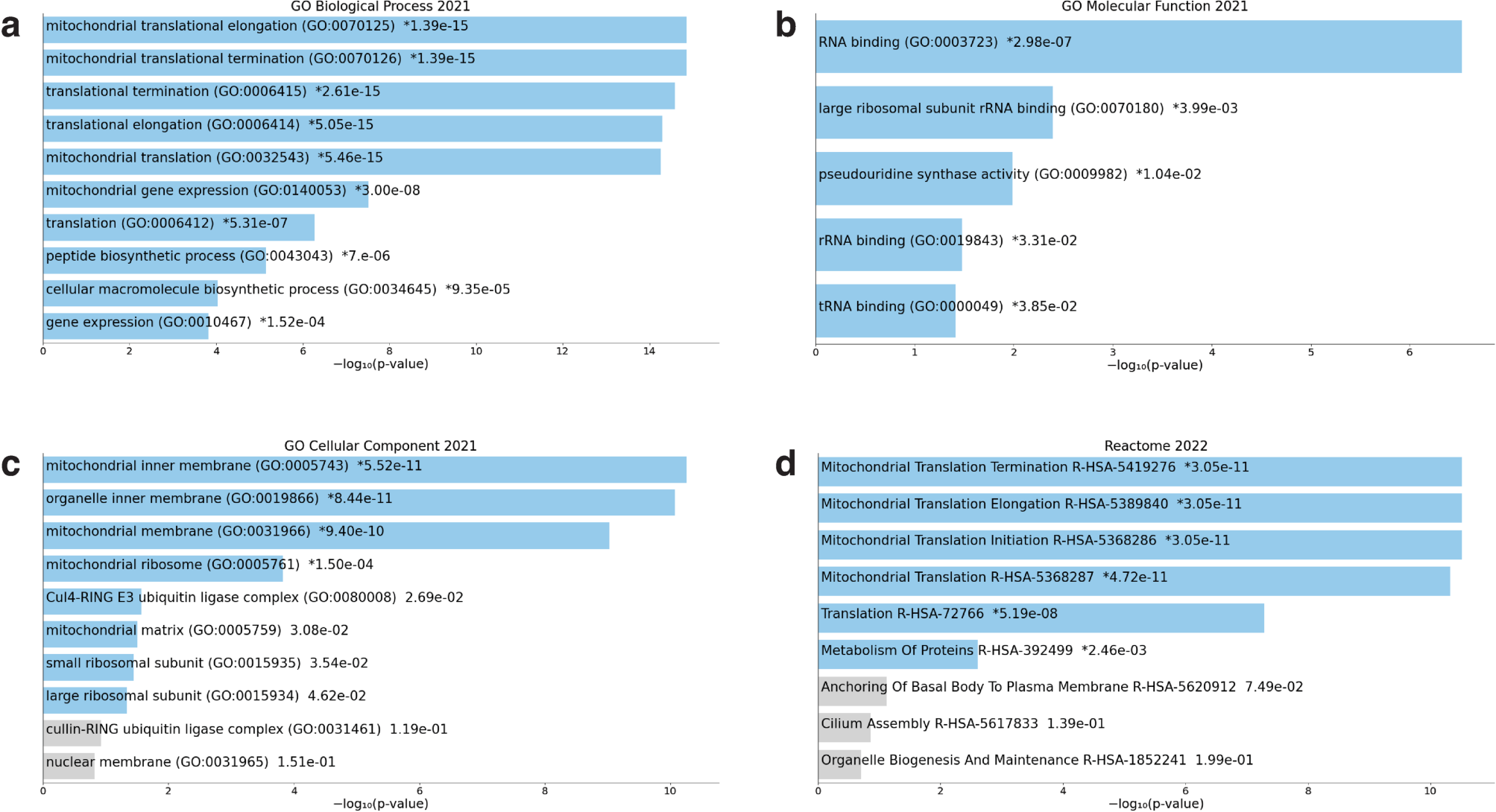
Gene ontology and pathway analysis bar charts. The term at the top has the most significant overlap with the input query gene set. The enriched terms for the hub dark genes are displayed based on log10(p-value), with the actual p-value shown next to each term. Coloured bars correspond to terms with significant p-values (< 0.05). An asterisk (*) next to a p-value indicates the term also has a significant adjusted p-value (< 0.05) (**a**) GO Biological Process (**b**) GO Molecular Function (**c**) GO Cellular Component (**d**) Reactome Pathways.

Overall, these results suggest that hub dark genes are closely linked to mitochondrial functions, particularly in the context of protein synthesis. This information can be invaluable for understanding the molecular mechanisms underlying various diseases and phenotypes associated with hub dark genes, as well as for potential therapeutic targeting in the future.

### Tissue expression of hub dark genes

We utilised the GTEx [45] database to investigate tissue-specific expression patterns of the 16 hub dark genes. Additionally, we searched for expression quantitative trait loci (eQTLs) within the dark genes to gain a better understanding of how genetic variants in these genes might contribute to disease development (see Methods section).

Our findings revealed that *RPUSD4* exhibited the highest median expression in muscle-skeletal (median TPM = 79.60). Similarly, *MRPL15* demonstrated the highest median expression in muscle-skeletal (median TPM = 109.8). Furthermore, *MRPL9* displayed the highest median expression in the uterus (median TPM = 77.75). *R3HDM2* highest median expression was observed in the brain-cerebellar (median TPM = 81.38), while *DCAF15* had the highest median expression within the testis (median TPM = 103.3 (Fig 11a).

**Fig 11.**
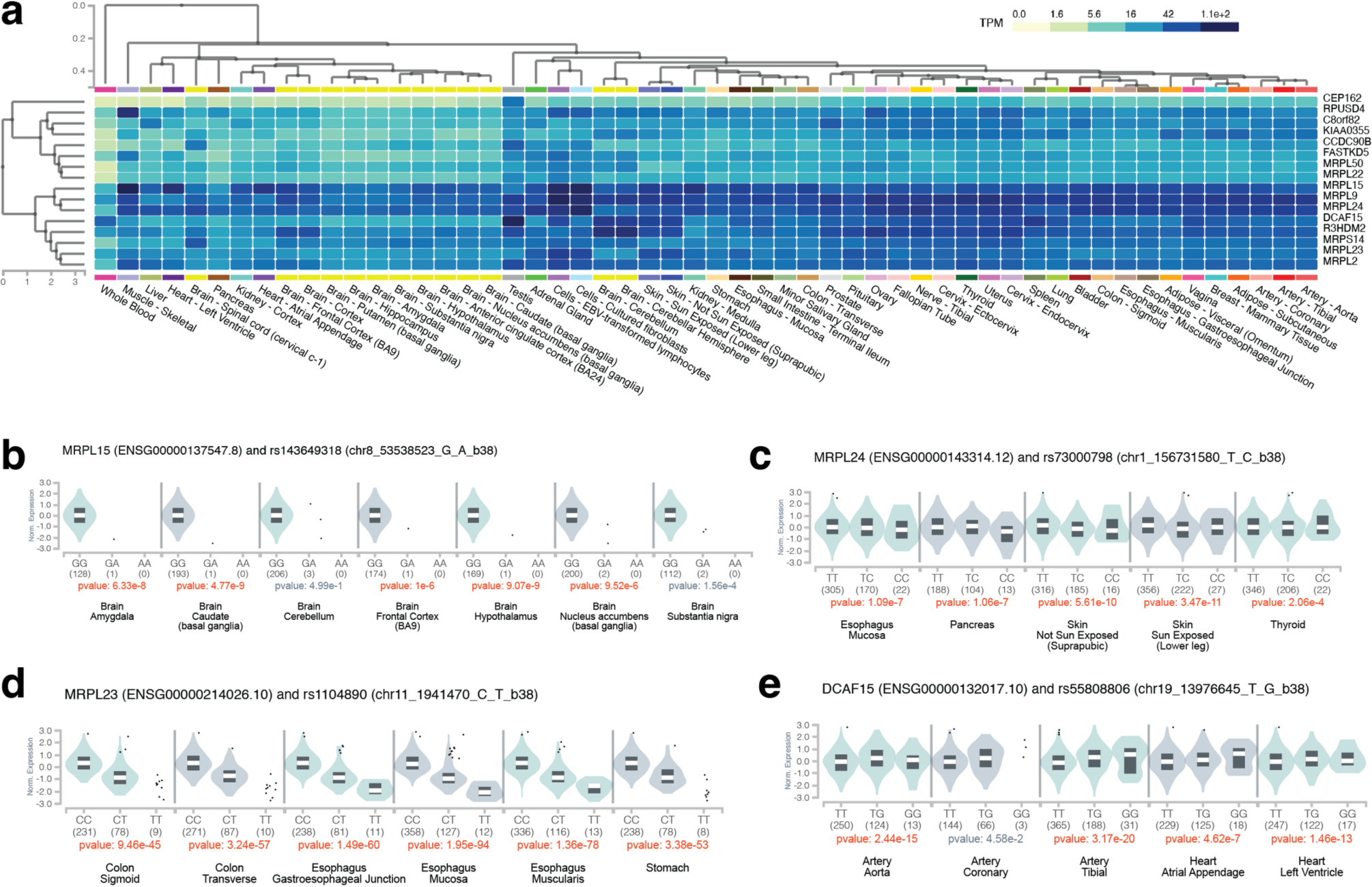
Tissue enrichments of Tdark hub genes. Expression of Tdark hub genes in various tissues **a.** and eQTLs and tissue expression of Tdark hub genes **b.** *MRPL15* **c.** *MRPL24* **d**. *MRPL23* **e.** *DCAF15*.

We also investigated the effects of eQTLs on the expression of hub dark genes in specific tissues. Interestingly, we found that 5,784 eQTLs significantly affected the expression of hub dark genes (p < 0.05) in various tissues, including the brain, oesophagus, and skin (see S5 File). For example, genetic variant rs143649318 (located at chr8_53538523_G_A_b38) significantly decreases the expression of *MRPL15* in the brain-amygdala (p = 6.33 x 10^-8^; Normalised effect size [NES] = -2.36), brain-hypothalamus (p = 9.07 x 10^-9^; NES = -2.19) and brain-frontal cortex (p = 1.00 x 10^-6^; NES = -1.99) (Fig 11b, also see S5 File). Furthermore, rs1104890 (located at chr11_1941470_C_T_b38) significantly decreases the expression of *MRPL23* in different tissues including the colon-transverse (p = 3.24 x 10^-57^; NES = -1.05), oesophagus-muscularis (p = 1.36 x 10^-78^; NES = -1.15), and stomach (p = 3.38 x 10^-53^; NES = -1.19) (Fig 11d, also see S5 File). Additionally, rs55808806 (located at chr19_13976645_T_G_b38) significantly increases the expression of *DCAF15* in the artery-aorta (p = 2.44 x 10^-15^; NES = 0.37); artery-tibial (p = 3.17 x 10^-20^; NES = 0.27) and heart-atrial appendage (p = 4.62 x 10^-7^; NES = 0.19) (Fig 11e, also see S5 File).

To further highlight the correlation between specific genetic variants and tissue-restricted phenotypes resulting from their impact on gene expression, we employed the Open Targets Genetics [46] web portal (https://genetics.opentargets.org) (see Methods section). Notably, we observed that rs1196456 (located at chr1_151764873_T_A_b38), which increases the expression of *MRPL9* in the heart (p = 1.10 x 10^-6^; NES = 0.23), exhibited an association with myocardial infarction (p = 1.80 x 10^-9^, Beta = 0.0578). Additionally, rs686646 (located at chr11_126189527_A_C_b38), which decreases *RPUSD4* expression in the pancreas (p = 7.90 x 10^-7^; NES = -0.2), displayed an association with diabetes (p = 4.20 x 10^-3^, Beta = 0.352). Similarly, rs1104890 (located at chr11_1941470_C_T_b38), which reduces *MRPL23* expression in skeletal muscle (p = 3.00 x 10^-119^; NES = -1.1), showed an association with intervertebral disc disorder (p = 7.60 x 10^-5^, Beta = 0.0569) (see S6 File for more details). Our findings demonstrate that understanding the tissue-specific expression patterns of genes is important for identifying genetic variants that contribute to disease susceptibility in specific tissues or organs.

### Tissue distribution and tissue specificity analysis of dark and light gene expression

We employed the Human Protein Atlas (HPA) [47] data and predictions from the Xpresso model [49] to conduct a comprehensive analysis of tissue distribution and tissue specificity in the context of dark and light gene expression. In terms of tissue distribution, our focus was specifically on categorising records based on the number of tissues in which mRNA transcripts were detected, namely, detected in all tissues, in many tissues, in some tissues, in a single tissue, or not detected in any tissue. For mRNA tissue specificity, we assessed records across different categories, including those that were group enriched, displayed low tissue specificity, were not detected, showed tissue enrichment, or exhibited tissue enhancement (see Methods section for the detailed description of categories). This approach allowed us to gain insights into the distinct tissue distribution and specificity of dark and light genes.

In the analysis of RNA tissue distribution, encompassing both dark and light genes, we examined 17,978 records. We found that the expression of mRNA transcripts from dark and light genes tends to vary across different tissues, with some transcripts being present in all tissues while others are limited to only a few. Notably, more than half of these records (9,154; 50.95%) revealed the presence of mRNA transcripts in all tissues. Meanwhile, a smaller fraction of the records demonstrated that mRNA transcripts were detected in many tissues (5,205; 28.98%) or some tissues (2,928; 16.29%) (Fig 12a and S7 File). This observation underscores the diverse tissue-specific expression patterns of dark and light genes.

**Fig 12.**
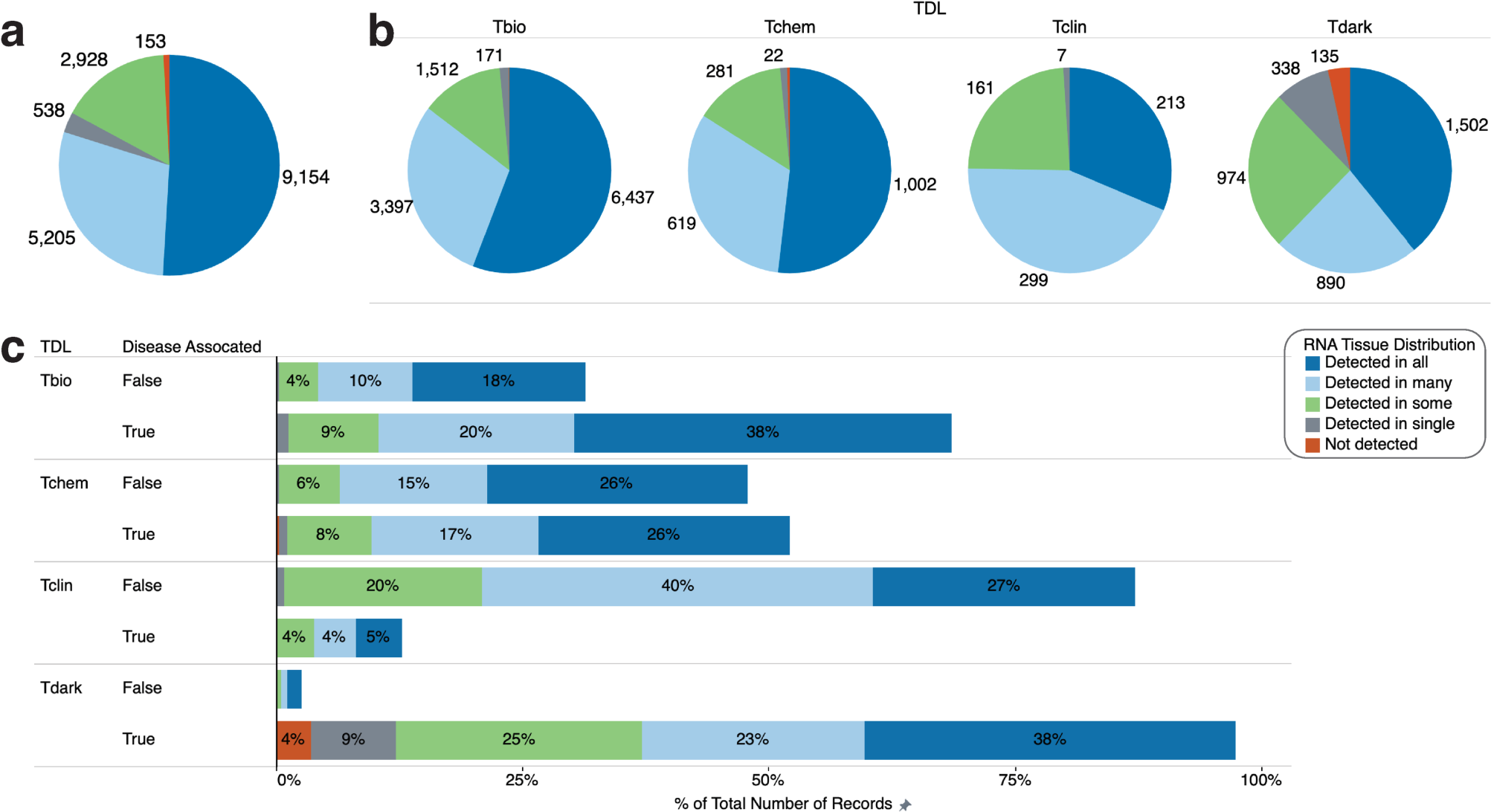
Tissue distribution of dark genes and light genes expression. **a.** The sum of the number of records on RNA tissue distribution. **b.** The sum of the number of records broken down by target development level (TDL). **c.** Percentage of the total number of records on RNA tissue distribution across distinct Target Development Level (TDL) classes linked to disease. The colour shows details about RNA tissue distribution. The size shows the sum of the number of records. The marks are labelled by the sum of the number of records.

Furthermore, the number of records broken down by TDL classes revealed that Tbio genes exhibited the highest number of RNA tissue distribution records detected in all tissues (6,437 records; 56%). Similarly, Tdark genes (1,502 records; 39%) and Tchem genes (1,002 records; 52%) also demonstrated the highest number of RNA tissue distribution records detected in all tissues. In contrast, Tclin genes displayed the highest number of RNA tissue distribution records detected in many tissues (299 records; 44%) (Fig 12b, also see S7a Fig). Additionally, we found that only a small percentage of records exhibited limited RNA tissue distribution, with 538 records (2.99%) detected in a single tissue and 153 records (0.85%) not detected in any tissue, suggesting restricted expression (Fig 12a). Notably, Tdark genes showed the highest percentage of such records, with 338 (63%) falling into the single-tissue detection category and 135 (88%) not being detected in any tissue, as depicted in Fig 12b and S7b Fig.

Furthermore, we observed variations in RNA tissue distribution across distinct TDL classes linked to disease. Our findings indicate that, among the TDL classes associated with disease, Tbio and Tdark genes demonstrated the highest prevalence of RNA tissue distribution records detected in all tissues (38% each), followed by Tchem genes (26%). Additionally, Tdark exhibited the highest number of RNA tissue distribution records detected in many tissues (23%), followed by Tbio (20%) and Tchem (17%). However, the proportion of genes identified in a single tissue was notably lower, ranging from 0% to 9%, with Tdark having the highest representation (Fig 12c). These results imply a substantial overlap in RNA tissue distribution across different tissues. Many genes are expressed in multiple tissues and may play a role in multiple disease pathways. However, some genes are tissue-specific and may play a more targeted role in disease development. Overall, these findings provide important insights into the molecular mechanisms underlying disease and highlight the importance of considering RNA tissue distribution in the development of targeted therapeutics.

In our analysis of RNA tissue specificity, encompassing both dark and light genes, we examined a total of 17,978 records. Our findings indicated that expression of these genes in most tissues (7,919 records; 44.05%) exhibited low tissue specificity, followed by tissue-enhanced (6,002 records; 33.39%) and tissue-enriched (2,431 records; 13.52%) categories (Fig 13a and S7 File). Furthermore, among the TDL classes, Tbio genes had the highest number of records with low tissue specificity (5,540 records; 48%), followed by tissue-enhanced and tissue-enriched categories [3,760 (33%) - tissue-enhanced; 1,327 (12%) - tissue-enriched]. Similarly, Tdark genes [1,418 (37%) - low tissue specificity; 1,237 (32%) - tissue-enhanced; 729 (19%) - tissue-enriched] and Tchem genes [803 (42%) - low tissue specificity; 716 (37%) - tissue-enhanced and 234 (12%) - tissue-enriched] displayed a similar pattern to the Tbio gene class. In contrast, Tclin genes had the highest number of records in the tissue-enhanced category, with 289 records (43%) (Fig 13b and S8a Fig).

**Fig 13.**
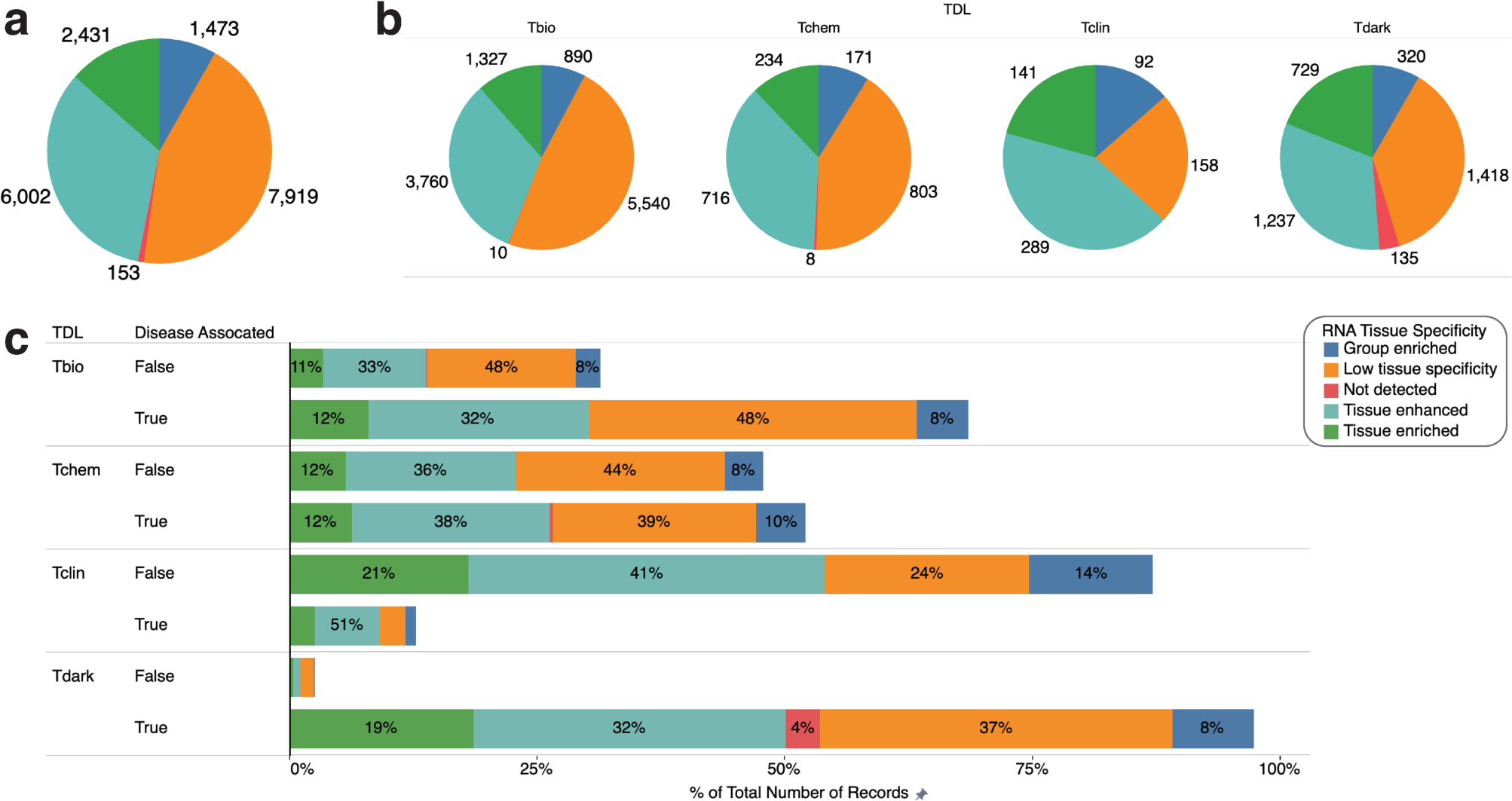
Tissue specificity of dark genes and light genes expression. **a.** The sum of the number of records on RNA tissue specificity. **b.** The sum of the number of records broken down by target development level (TDL). **c.** Percentage of the total number of records on RNA tissue specificity across distinct Target Development Level (TDL) classes linked to disease. The colour shows details about RNA tissue specificity. The size shows the sum of the number of records. The marks are labelled by the sum of the number of records.

Overall, we found that the total number of records for each category of RNA tissue specificity broken down by TDL classes was consistently high in Tbio, followed by Tdark, Tchem and lastly, Tclin, except for the “Not detected” category, which had Tdark with the highest record (135 records; 88%) (Fig 13b and S8b Fig). Additionally, low tissue specificity records remained consistently high in Tbio, Tchem, and Tdark, suggesting that these genes may have more generalised functions that are not confined to specific biological contexts, potentially playing crucial roles in multiple tissues or biological processes.

Moreover, we have identified distinctions in RNA tissue specificity among diverse TDL classes linked to disease. Notably, Tclin exhibited the highest share of genes displaying tissue enhancement (51%). Furthermore, our analysis revealed that among Tdark genes, 37% displayed low tissue specificity, 32% showed tissue enhancement, 19% were tissue-enriched, 8% were group-enriched, and 4% were not detected (Fig 13c). These findings suggest that a significant portion of Tdark genes is expressed in particular tissues or groups of tissues, implying potential involvement in specific physiological processes. However, a considerable number of Tdark genes exhibited low tissue specificity, suggesting that their functions may be less tissue-specific or not yet fully understood. These findings underscore the significance of further exploration of the role of Tdark genes in disease development and tissue-specific processes.

Furthermore, our investigation has yielded noteworthy insights into the RNA tissue distribution and tissue specificity of dark and light genes. Notably, mRNA transcripts of both dark and light genes detected in all tissues exhibited the highest number of records with low levels of tissue specificity, with Tbio genes being the most prominent (55%), followed by Tdark genes (13%). While a subset of records demonstrated tissue enhancement for mRNA transcripts detected in many tissues, with the highest proportion in Tbio genes (39%), followed by Tdark genes (11%) and Tchem genes (7%). On the other hand, mRNA transcripts detected in single tissues, categorised as tissue enriched, had the highest proportion in Tdark genes (45%), followed by Tbio genes (29%). These findings contribute to our understanding of gene expression regulation and provide insights into the distinct functions associated with specific tissues (see S9 Fig).

Overall, these findings collectively underscore the importance of exploring the distinct roles of Tdark genes in disease development and tissue-specific processes. While there are similarities, there are notable differences in the RNA tissue distribution and RNA tissue specificity between dark and light genes, emphasising their unique contributions to biological functions and disease pathways.

## Discussion

We have conducted integrative analyses to illuminate the potential roles of Tdark genes and their possible contribution to genetic diseases and phenotypes. Our findings reveal that the number of genes associated with genetic traits varies depending on the target development level. Tdark genes display a relatively high average count of gene associations per disease, indicating the complex or multifaceted roles these dark genes may play in disease pathogenesis. Additionally, Tdark genes have the lowest number of diseases per gene, possibly due to limited research or understanding [23,25]. Further research is required to understand the function and potential significance of Tdark genes in various diseases and phenotypes. Furthermore, our analysis highlights psoriasis as the most prevalent disease across all development levels, emphasising the importance of studying Tdark genes in relation to this condition.

We identified certain transcription factors as potential regulators of both light and dark genes associated with specific genetic diseases. For instance, SUZ12 and TP63 were identified as potential regulators of dark and light gene expression in interstitial cystitis, while UBTF and NFE2L2 were identified as potential regulators of both dark and light genes associated with tuberculosis. These transcription factors have been implicated in various biological processes [62–66], and their identification in both light and dark genes suggests their important role in the pathogenesis of genetic diseases and phenotypes.

Our analysis of the gene-disease network (GDN) revealed the presence of connections among genetic diseases and phenotypes. Consistent with prior research [55], we found that out of 557 genetic traits examined, 503 belonged to a giant component, suggesting shared genetic origins to some extent [55,67]. Furthermore, there was considerable variation in the number of genes associated with different diseases and phenotypes, ranging from just a few to dozens, as seen in conditions like psoriasis, pick disease, tuberculosis, ulcerative colitis, interstitial cystitis, and Crohn’s disease. This variation may suggest the presence of shared genetic pathways or biological mechanisms [68–71]. The connections among disease genes in the disease-gene network (DGN) indicate their level of phenotypic similarity and involvement in shared disease pathways. Integrating this network with other interaction types (e.g., protein-protein interactions, transcription factor-promoter interactions, and metabolic reactions) can unveil novel genetic interactions and provide a more comprehensive understanding of disease mechanisms [55]. Notably, *CALHM6, HCP5, PRRG4, DDX60L,* and *RASA2* were identified as major hubs in the network, indicating their key roles in disease pathways [72,73]. Additionally, *PLBD1, CTRB2, C6orf15, DPM3,* and *CALHM6* exhibited high betweenness centrality, signifying their importance in regulating information flow among genes in the network [56,74].

We identified 16 most significant hub dark genes from the PPI network including *RPUSD4*, *FASTKD5, MRPL15, MRPL9, MRPL22, MRPL2, MRPL50, MRPL24, MRPL23, DCAF15, C8orf82, KIAA0355, CCDC90B, MRPS14, R3HDM2,* and *CEP162*. The interactions among most of these genes reinforces their potential functional significance, offering clues about their biological roles [75], and suggesting their possible involvement in the same biological pathways. Upon further analysis of hub genes through Enrichr, we found that the hub genes are significantly associated with GO (Gene Ontology) biological processes related to mitochondrial translation, such as translational elongation and termination. Therefore, we suggest that these genes may regulate and maintain mitochondrial protein synthesis, which is essential for cellular function [76,77]. The hub genes are enriched in GO cellular components like mitochondrial inner membrane, and GO molecular functions include RNA binding and large ribosomal subunit rRNA binding. Additionally, the hub dark genes participate in mitochondrial translation pathways, encompassing initiation, elongation, and termination processes. This finding indicates that hub genes are crucial in preserving the structural and functional integrity of subcellular compartments and in overseeing gene expression and protein synthesis within cells. Recent research has underscored the significant role of these hub dark genes in disease, with studies indicating their involvement in various pathologies and potential as therapeutic targets [78–81].

We conducted a detailed analysis of the expression patterns of 16 hub dark genes across diverse tissues. The findings reveal that these genes are expressed in multiple tissues, suggesting their crucial role in a wide range of biological processes. For example, we found that *RPUSD4* is highly expressed in muscle-skeletal tissues, whereas *MRPL9* is expressed in EBV-transformed lymphocytes, cultured fibroblasts, and the uterus. In addition, *MRPL15* is expressed in muscle-skeletal and heart-left ventricles. Interestingly, the expression of quantitative trait loci (eQTLs) significantly influenced the expression of hub dark genes across various tissues, indicating that these genetic variants are instrumental in modulating the expression of hub dark genes in diverse biological contexts [82]. The discovery that specific eQTLs, such as rs1104890 and rs143649318, have a significant impact on the expression of specific hub dark genes in different tissues highlights the importance of understanding the tissue-specific regulation of gene expression [83]. Furthermore, these eQTLs provide potential markers for further investigation into the mechanisms underlying the regulation of hub dark genes [84]. Investigating the impact of these eQTLs on gene expression and protein function can provide insights into the molecular pathways regulating hub dark genes and potentially identify therapeutic targets for diseases associated with these genes [85–87].

In conclusion, our research demonstrates that dark genes are crucial in the pathogenesis of genetic diseases and phenotypes, revealing their multifaceted involvement in disease mechanisms. Moreover, network analysis exposes a common genetic foundation across various diseases, with certain genes acting as key hubs within these networks. Importantly, these hub dark genes are frequently involved in essential mitochondrial functions. The distinct expression patterns of these genes and the modulation by eQTLs emphasise the complexity of gene regulation in different biological contexts. Exploring dark genes further can enhance our understanding of genetic diseases and lead to the discovery of new therapeutic avenues.

## Data availability

All relevant data are within the manuscript and its supporting information files.

## Supporting information

Description of Supplementary Files

## Author contributions

**Conceptualisation:** Doris Kafita, Kevin Dzobo and Musalula Sinkala.

**Methodology:** Doris Kafita, Panji Nkhoma, Kevin Dzobo and Musalula Sinkala.

**Formal analysis:** Doris Kafita, Panji Nkhoma, Kevin Dzobo and Musalula Sinkala.

**Visualisation:** Doris Kafita, Kevin Dzobo and Musalula Sinkala.

**Writing – original draft:** Doris Kafita and Musalula Sinkala.

**Writing – review & editing:** Doris Kafita, Panji Nkhoma, Kevin Dzobo and Musalula Sinkala.

