## Supplementary material for "Unveiling the role of Tdark genes in genetic diseases and phenotypes through bioinformatics-based functional enrichment and network analyses": Description of Supplementary Files

### Supplementary Figures

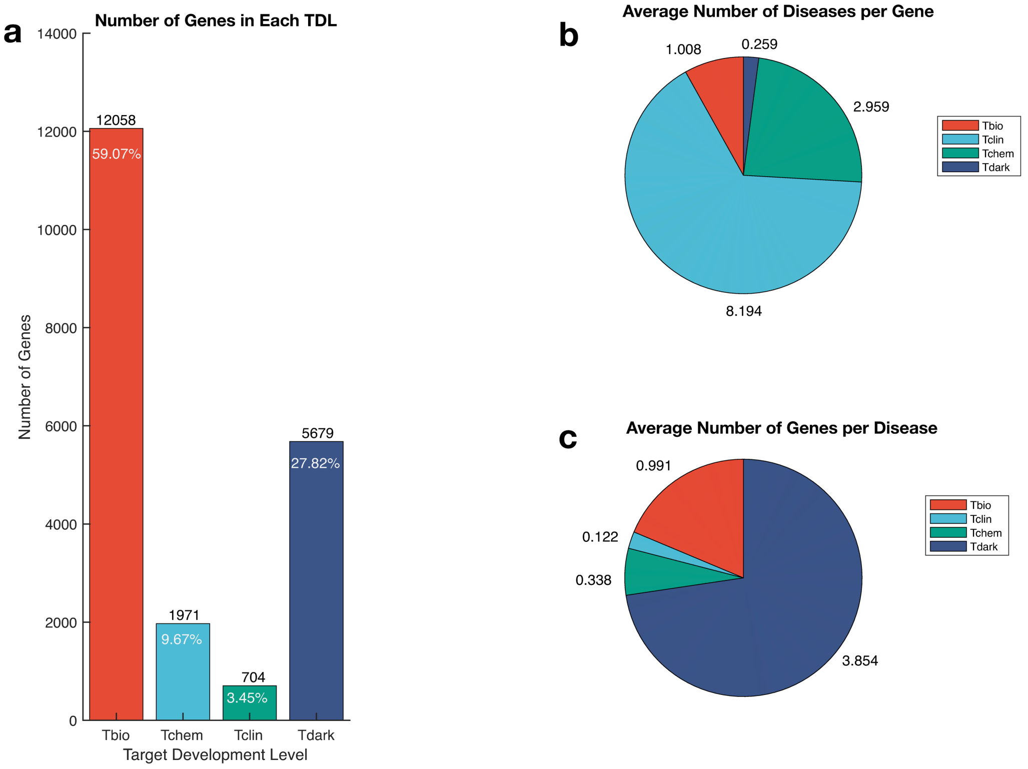


**S1 Fig. Distribution of genes associated with genetic diseases/phenotypes in each target development level (TDL). a**. Frequency of genes in each TDL in Pharos. **b.** Average number of diseases per gene **c.** Average number of genes per disease.

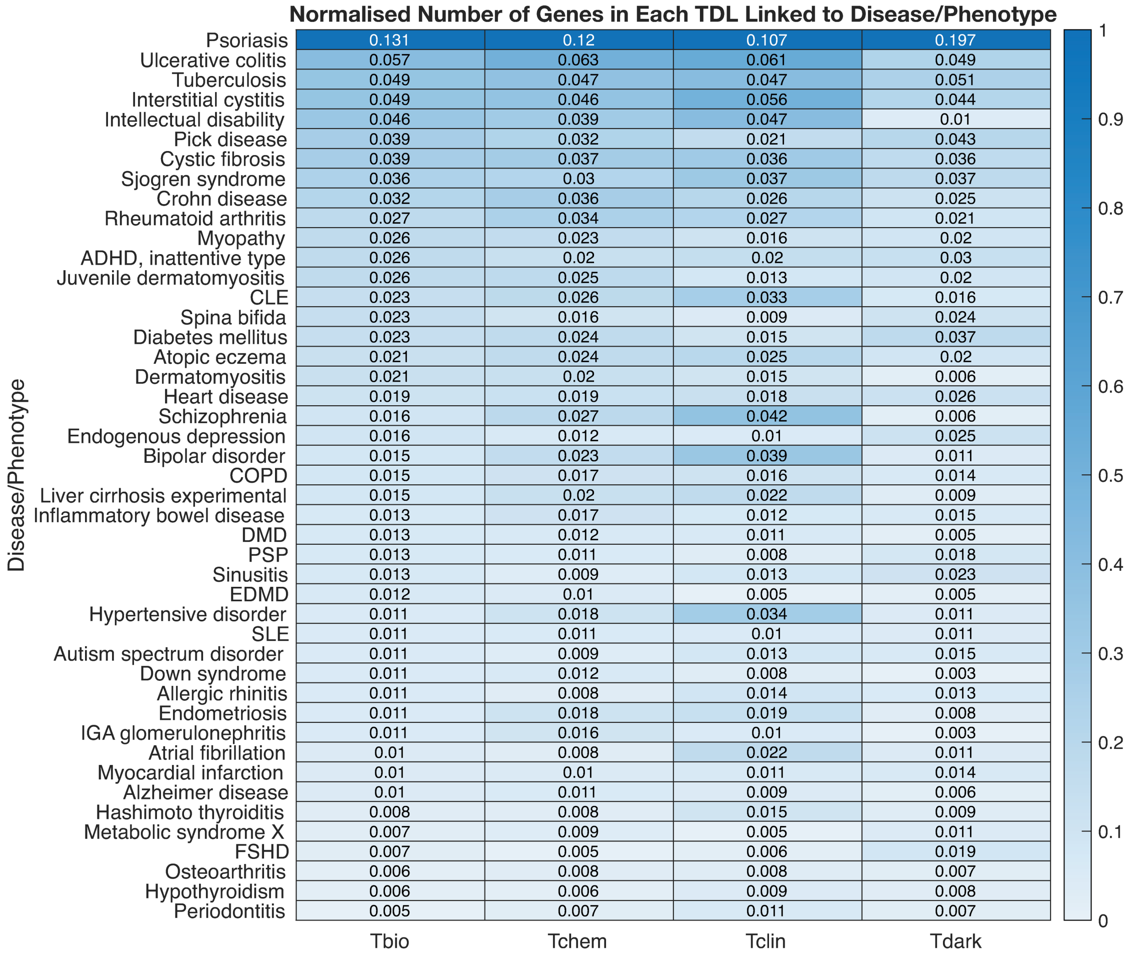


**S2 Fig. Normalised frequency of genes linked to the top 40 genetic diseases and phenotypes.** CLE: Cutaneous lupus erythematosus, COPD: Chronic obstructive pulmonary disease, DMD: Duchenne muscular dystrophy, PSP: Progressive supranuclear palsy, EDMD: Emery-Dreifuss muscular dystrophy, SLE: Systemic lupus erythematosus, FSHD: Facioscapulohumeral muscular dystrophy.

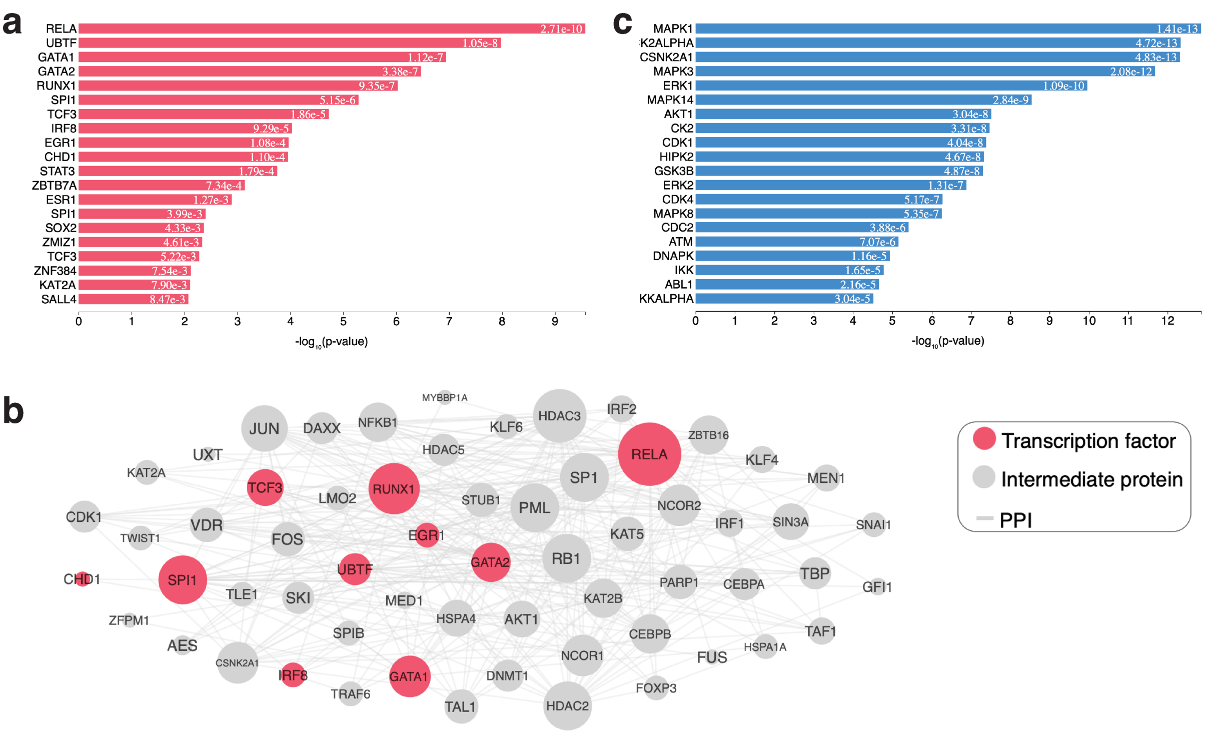


**S3 Fig. ChEA and KEA analyses of light genes associated with rheumatoid arthritis. a**. The top-20 predicted regulatory transcription factors. **b.** A subnetwork of connected transcription factors and their interacting proteins: the sub-network has 61 nodes with 669 edges. Transcription factors are the pink nodes, whereas the proteins that connect them are in grey. **c.** The top-20 predicted regulatory kinases.

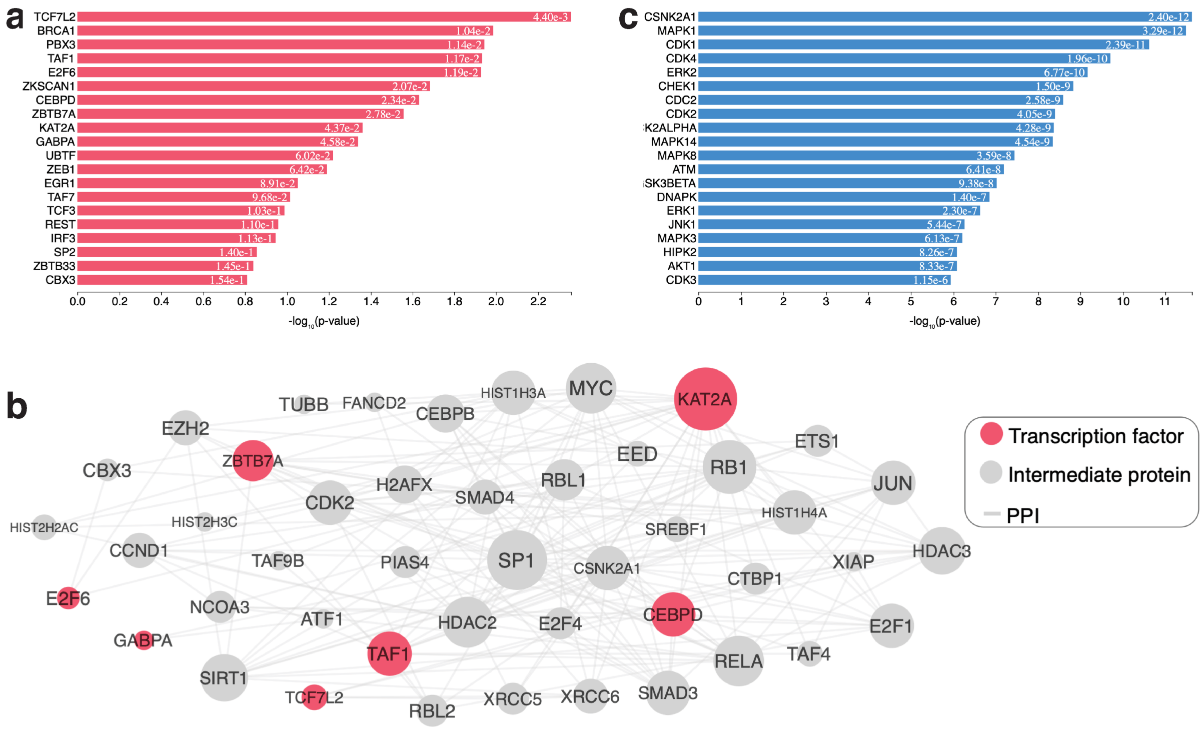


**S4 Fig. ChEA and KEA analyses of dark genes associated with pick disease. a.** The top-20 predicted regulatory transcription factors. **b.** A subnetwork of connected transcription factors and their interacting proteins: the sub-network has 49 nodes with 374 edges. Transcription factors are the pink nodes, whereas the proteins that connect them are in grey. **c.** The top-20 predicted regulatory kinases.

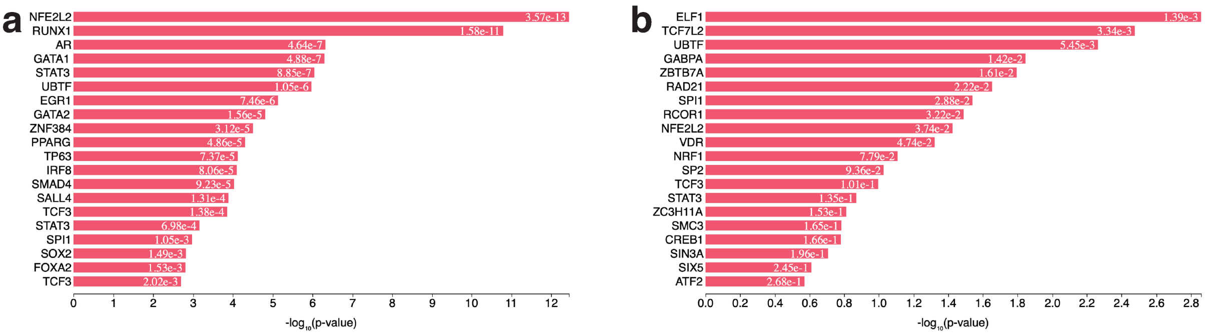


**S5 Fig. ChEA analysis of light and dark genes associated with tuberculosis. a.** The top-20 predicted regulatory transcription factors for light genes. **b.** The top-20 predicted regulatory transcription factors for dark genes.

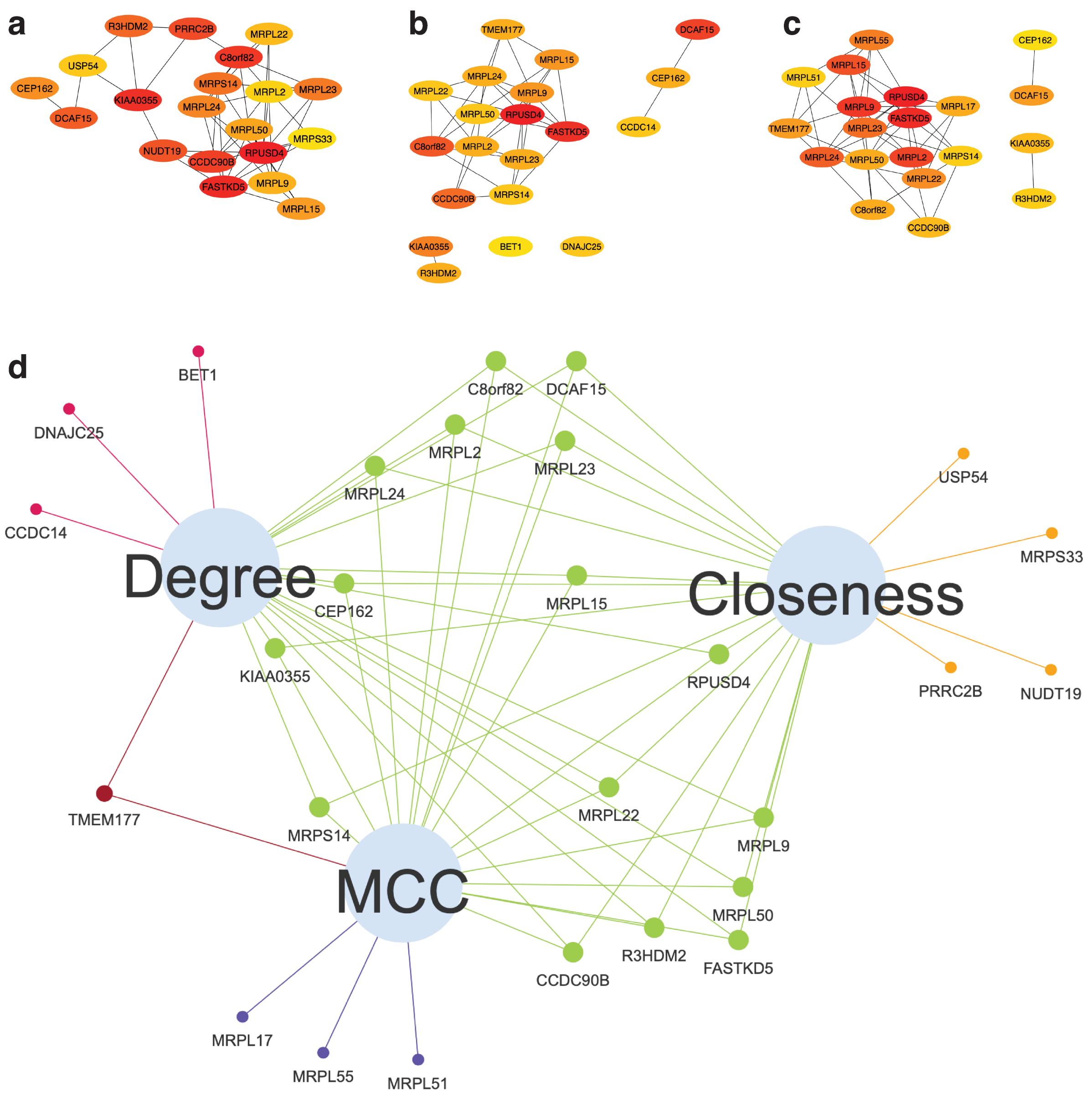


**S6 Fig. Hub genes identified by 3 different algorithms. a.** Closeness centrality. **b.** Degree centrality and **c.** Maximal Clique Centrality (MCC). **d.** The Venn network, determined by functional protein association PPI network analysis, reveals 16 overlapping hub genes. Nodes highlighted in green indicate the common hub genes identified by all three algorithms.

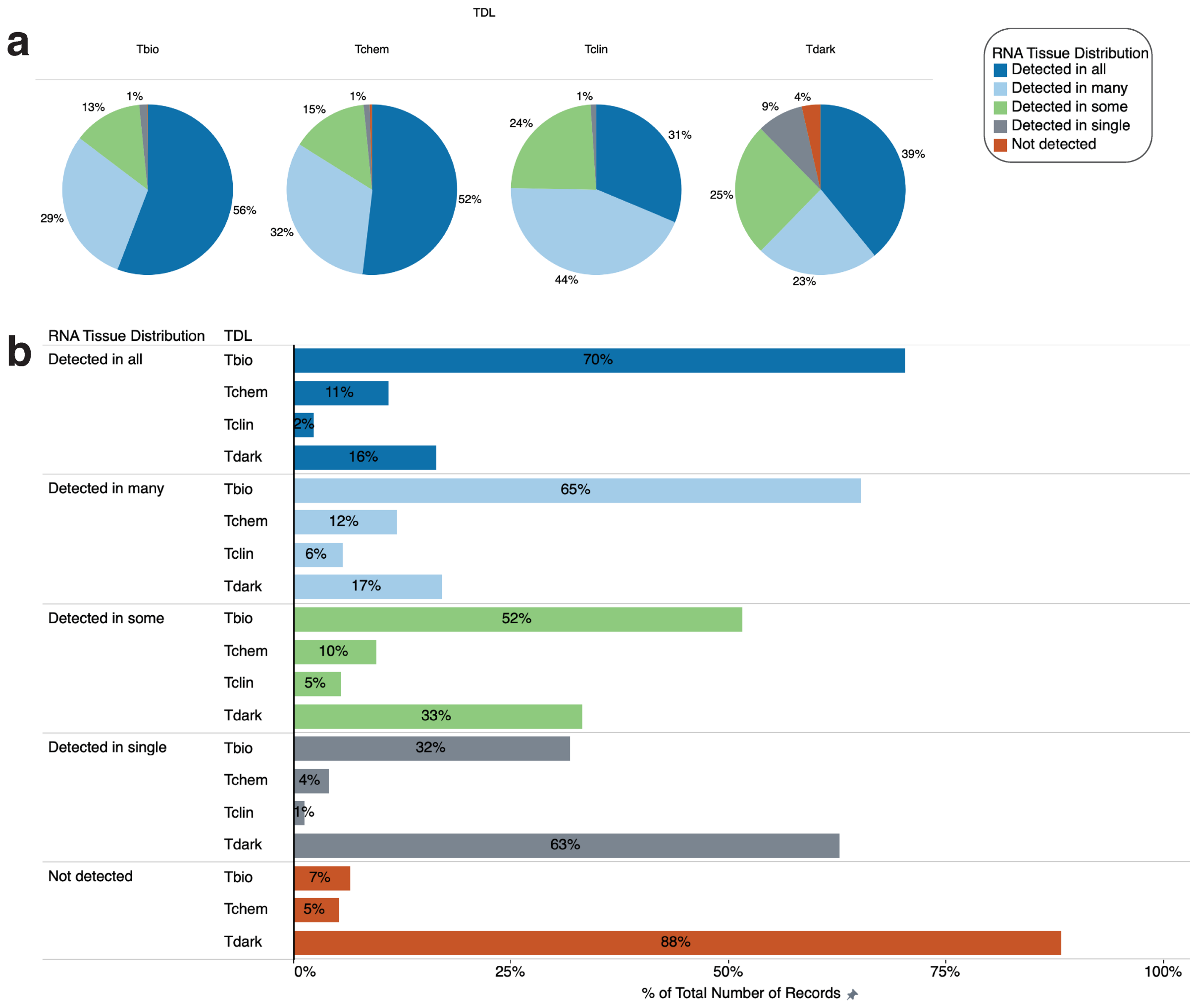


**S7 Fig. Tissue distribution. a.** Percentage of the number of records broken down by target development level (TDL). **b.** Percentage of total number of records for each TDL broken down by RNA tissue distribution. The colour shows details about RNA tissue distribution, with the marks labelled by the percentage of the total number of records.

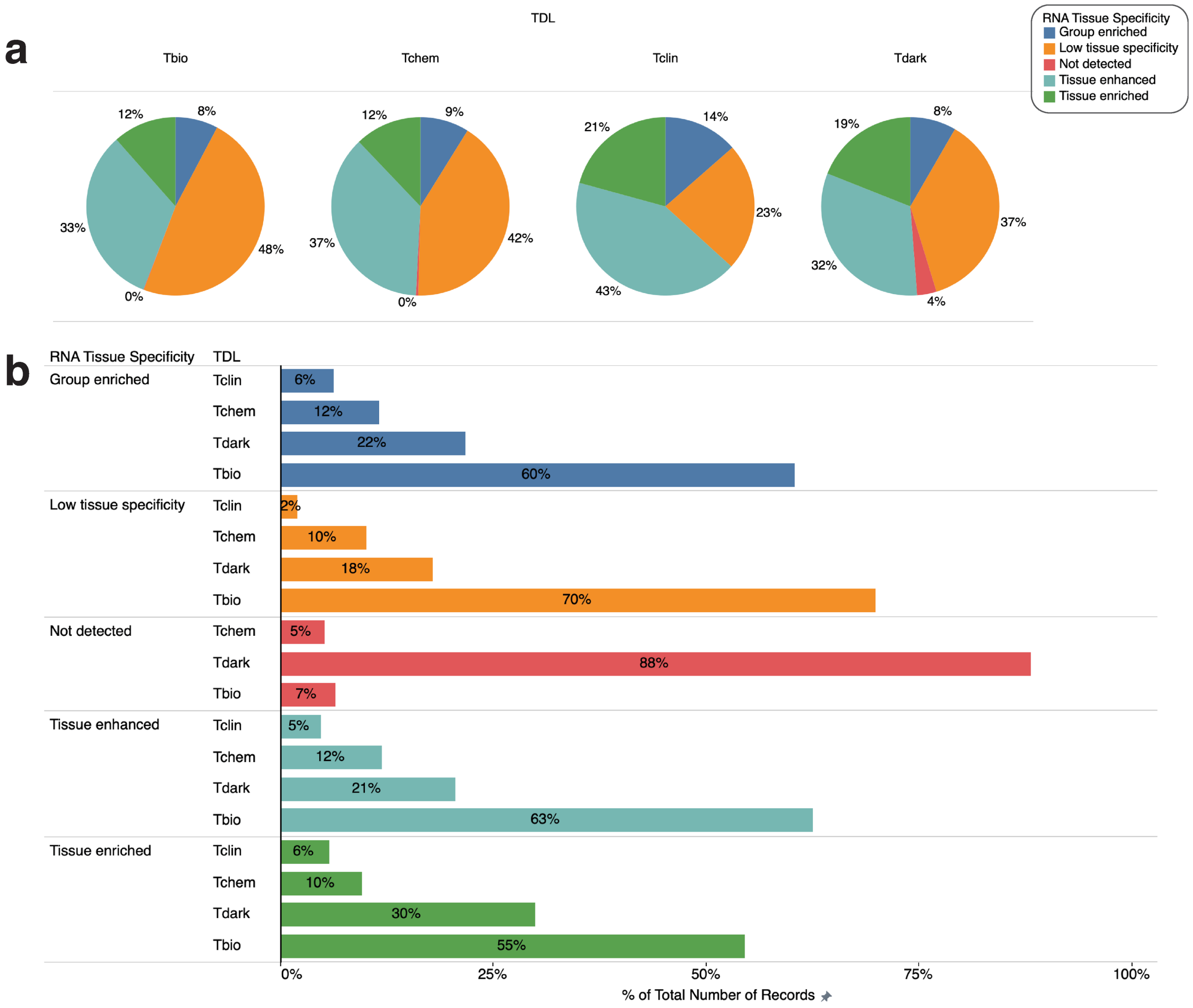


**S8 Fig. Tissue specificity. a.** Percentage of the number of records broken down by target development level (TDL). **b.** Percentage of total number of records for each TDL broken down by RNA tissue specificity. The colour shows details about RNA tissue specificity, with the marks labelled by percentage of the total number of records.

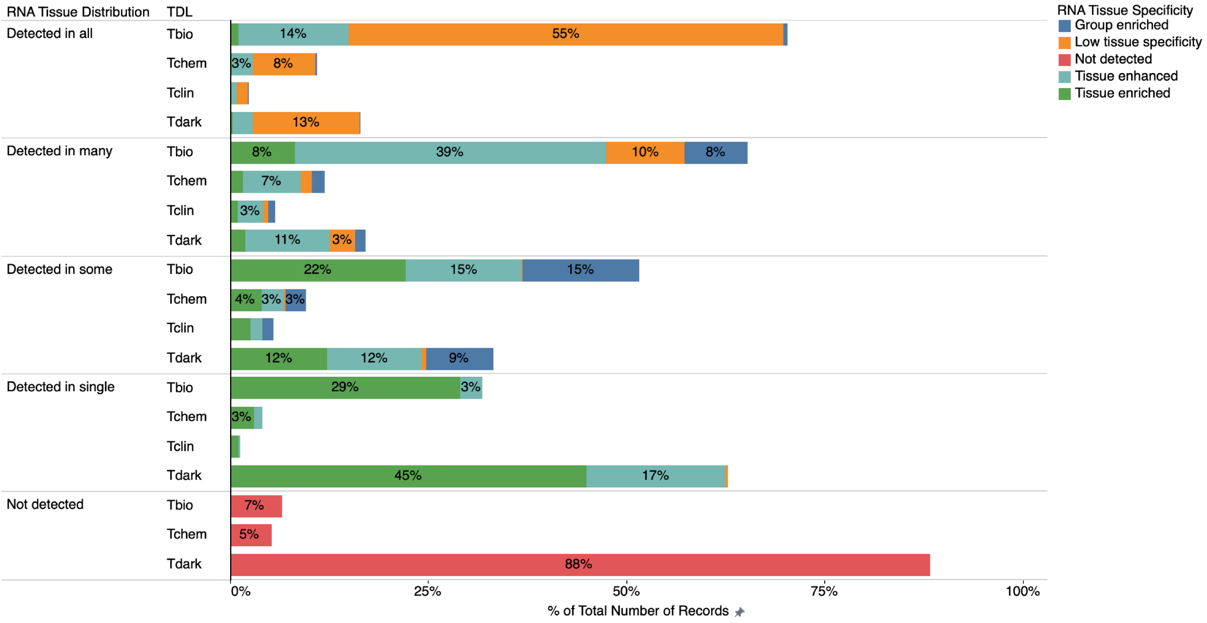


**S9 Fig. Tissue distribution and tissue specificity analysis of dark genes and light genes expression.** Percentage of the total number of records for each target development level (TDL) broken down by RNA tissue distribution. The colour shows details about RNA tissue specificity, with the marks labelled by percentage of the total number of records.
