## Supplementary Figures for "Unveiling the role of Tdark genes in genetic diseases and phenotypes through bioinformatics-based functional enrichment and network analyses"

### Description of Additional Supplementary Files

**S1 File. Supplementary data of light and dark genes and genetic diseases/phenotypes.** The spreadsheet contains the following results/datasets according to the sheet name. *Pharos data:* Distribution of genes and genetic diseases/phenotypes within each target development level obtained from Pharos. *Tdark dxsphenotypes:* Frequency of genetic diseases and phenotypes associated with Tdark genes. *Tbio dxsphenotypes:* Frequency of genetic disease and phenotypes associated with Tbio genes. *Tchem dxsphenotypes:* Frequency of genetic disease and phenotypes associated with Tchem genes. *Tclin dxsphenotypes:* Frequency of genetic disease and phenotypes associated with Tclin genes. *Dxs-PhenotypesLinkedtoTdarkOnly:* List of genetic diseases and phenotypes exclusively associated with Tdark genes.

**S2 File. Transcription factor and kinase enrichment analyses of dark and light genes associated with the top 10 genetic diseases and phenotypes.** The spreadsheet contains predicted regulatory transcription factors and kinases. The sheet names are structured based on the category of genes (dark or light genes), the type of analysis and the specific genetic disease e.g., *Dark CHEA-Tuberculosis*: contains predicted transcription factors potentially regulating the expression of dark genes linked to tuberculosis, along with a list of regulated genes from the input gene list. *Dark KEA-Tuberculosis;* contains a list of protein kinases that likely regulate the transcriptome signature of tuberculosis.

**S3 File. Network analysis of dark genes and the associated diseases/phenotypes.** The spreadsheet contains the following results/datasets according to the sheet name. *DiseasenetworkGDN:* Weighted degree centrality, betweenness centrality, closeness centrality and eccentricity for the gene-disease network (GDN). *GenenetworkDGN:* Weighted degree centrality, betweenness centrality, closeness centrality and eccentricity for the disease-gene network (DGN). The remaining sheets contain results for the seven smaller network clusters within the diseasome bipartite, including the following metrics: degree, eccentricity, closeness centrality and betweenness centrality.

**S4 File. Function enrichment of hub dark genes.** The spreadsheet contains the following results according to the sheet name. *GO Biological:* Gene ontology for biological processes. *GO Cellular:* Gene ontology for cellular components. *GO Molecular:* Gene ontology for molecular functions. *Pathways-Reactome:* Reactome pathway analysis results.

**S5 File. Tissue expression of hub dark genes.** The spreadsheet contains a list of expression quantitative trait loci (eQTLs) that significantly impact the expression of hub dark genes in various tissues.

**S6 File. Correlation between specific genetic variants and tissue-restricted phenotypes.** The spreadsheet contains the following results according to the sheet name. *MRPL9 1_151764873_T_A-associate:* Tissue restricted phenotypes for rs1196456. *MRPL23 11_1941470_C_T-associate:* Tissue restricted phenotypes for rs1104890. *RPUSD4 11_126189527_A_C-associa:* Tissue restricted phenotypes for rs686646.

**S7 File. Tissue distribution and tissue specificity of dark and light gene expression.** The spreadsheet contains the following results/datasets according to the sheet name. *Tissue distribution:* mRNA Tissue distribution of dark and light genes Tissue specificity: mRNA tissue specificity of dark and light genes.
